# Epigenetic repression of cFos supports sequential formation of distinct spatial memories

**DOI:** 10.1101/2024.02.16.580703

**Authors:** Andreas Franzelin, Paul J. Lamothe-Molina, Christine E. Gee, Andrey Formozov, Eric R. Schreiter, Fabio Morellini, Thomas G. Oertner

## Abstract

Expression of the immediate early gene cFos modifies the epigenetic landscape of activated neurons with downstream effects on synaptic plasticity. The production of cFos is inhibited by a long-lived isoform of another Fos family gene, ΔFosB. It has been speculated that this negative feedback mechanism may be critical for protecting episodic memories from being overwritten by new information. Here, we investigate the influence of ΔFosB inhibition on cFos expression and memory. Hippocampal neurons in slice culture produce more cFos on the first day of stimulation compared to identical stimulation on the following day. This downregulation affects all hippocampal subfields and requires histone deacetylation. Overexpression of ΔFosB in individual pyramidal neurons effectively suppresses cFos, indicating that accumulation of ΔFosB is the causal mechanism. Water maze training of mice over several days leads to accumulation of ΔFosB in granule cells of the dentate gyrus, but not in CA3 and CA1. Because the dentate gyrus is thought to support pattern separation and cognitive flexibility, we hypothesized that inhibiting the expression of ΔFosB would affect reversal learning, i.e., the ability to successively learn new platform locations in the water maze. The results indicate that pharmacological HDAC inhibition, which prevents cFos repression, impairs reversal learning, while learning and memory of the initial platform location remain unaffected. Our study supports the hypothesis that epigenetic mechanisms tightly regulate cFos expression in individual granule cells to orchestrate the formation of time-stamped memories.

## Introduction

Current models of memory storage suggest that memory retrieval requires the reactivation of a subset of neurons that were active during the lived experience. Fear conditioning experiments in TetTag mice provide compelling experimental evidence for this concept (Liu et al. 2012; Reijmers et al. 2007). In these mice, cFos activation is used to label the most active neurons within a time window set by doxycycline withdrawal. cFos-expressing neurons (cFos^+^) in the hippocampus appear to play a distinct role in the network. Calcium imaging in cFos reporter mice showed that cFos^+^ neurons have very stable place fields and form ensembles with significantly correlated activity (Pettit et al. 2022). The goal of this study was to better understand the conditions for cFos activation, particularly the rapidly changing expression patterns seen in the dentate gyrus (DG) (Lamothe-Molina et al. 2022). It is well established that action potential firing leads to a proportional increase in intracellular calcium concentration ([Ca^2+^]_i_) (Maravall et al. 2000). Calcium release from the endoplasmic reticulum produces even larger [Ca^2+^]_i_ transients (Holbro, Grunditz, and Oertner 2009; Mellentin, Jahnsen, and Abraham 2007). Notably, nuclear [Ca^2+^]_i_ is a potent trigger for epigenetic modifications (Bading 2013; Mozolewski et al. 2021), making it a compelling parameter to study in the context of immediate early genes (Bengtson et al. 2010).

To examine the correlation between activity, nuclear calcium and cFos expression, we used organotypic slice cultures of the rat hippocampus. This culture system effectively preserves neuronal identities and connectivity between the different hippocampal subfields. To simultaneously record nuclear calcium in all neurons, we used CaMPARI2 - a recently improved calcium integrator (Moeyaert et al. 2018). CaMPARI2 contains a green fluorescent protein (mEOS), which irreversibly converts to a red fluorescent form when bound to calcium and exposed to violet illumination (405 nm). To measure calcium where it is most important for epigenetics, we used a nuclear-localized version, H2B-CaMPARI2. An additional advantage of nuclear targeting was the easy detection and segmentation of fluorescent nuclei with commercial image analysis software, a task that is challenging to automate when using cytoplasmic indicators. To obtain a permanent cell-specific record of activity, we illuminated the cultures with violet light inside the incubator during the stimulation period. Photoconversion of individual nuclei was quantified after fixation on a confocal microscope. This process allowed us to evaluate calcium transients that occurred one hour before fixation within a defined 60-second window.

Our experiments with slice cultures indicated that ΔFosB accumulation induces an epigenetic mechanism that prevents repeated expression of cFos in individual neurons. To determine whether this mechanism is actually evoked during learning, we tested whether inhibiting histone modification would affect performance in a complex spatial learning task (Morellini 2013). Indeed, we found that mice had no problem learning a single platform position under HDAC inhibition, but performed much worse when challenged with daily changing positions. Our findings support the concept that epigenetic mechanisms control the dynamic allocation of “fresh” neurons for memory storage, resulting in well-separated episodic memories (Watrous and Ekstrom 2014; Chowdhury and Caroni 2018).

## Results

### Nuclear calcium does not predict cFos expression level

To investigate the relationship between activity-induced increases in nuclear [Ca^2+^] and cFos expression in many neurons in parallel, we virally transduced neurons with the photoconvertible calcium indicator CaMPARI2 fused to histone H2B for nuclear targeting (H2B-CaMPARI2). When bound to calcium, illumination of CaMPARI2 at 405 nm induces an irreversible conversion from a green to a red state, allowing calcium levels to be integrated in a precise time window defined by the photoconversion light. After five days of H2B-CaMPARI2 expression, we stimulated slice cultures with a drop of the GABA_A_ antagonist bicuculline (Fig. 1A), which induces bursts of synchronized activity (Fig. S1). After 60 s of bicuculline treatment, the induced activity was terminated by pipetting the Na^+^ channel blocker tetrodotoxin (TTX, 300 µl, 1 µM) onto the slice culture. During the stimulation period, the slice cultures were continuously illuminated (395 nm) to ensure photoconversion of calcium-bound H2B-CaMPARI2. Cultures were fixed 60 min after the end of the stimulation period and immunostained for cFos and the red-converted version of CaMPARI2. At a viral titer of 10^12^ vg/ml, cFos expression in H2B-CaMPARI2-expressing cultures was not different from non-transfected control cultures (Fig. S2). As negative controls, we included cultures that were treated with TTX instead of bicuculline (Fig. S3).

**Figure 1:**
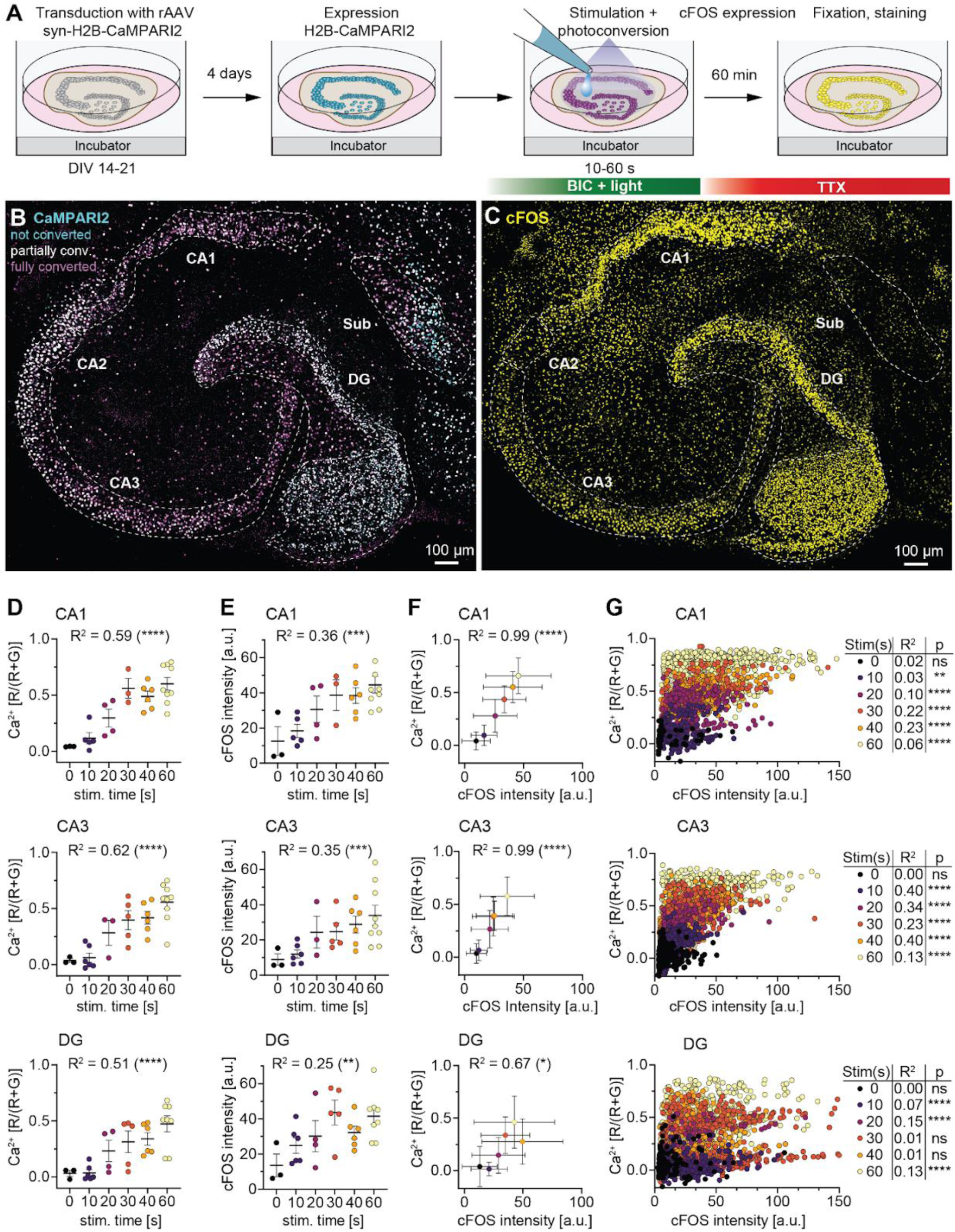
Correlations between nuclear calcium levels and cFos expression. **A)** Time line of in-incubator stimulation experiments using synapsin-H2B-CaMPARI2-transduced slice cultures, stimulated with bicuculline (BIC) followed by tetrodotoxin (TTX). **B)** Slice culture stimulated for 60 s with bicuculline, showing strong CaMPARI2 photoconversion (cyan = non-converted, white = partially converted, magenta = fully converted). **C)** Anti-cFos staining (yellow), same tissue as in B. **D)** CaMPARI2 photoconversion (Ca^2+^, [R\(R+G)]) as a function of stimulation time. Circles represent averages from individual slice cultures (40 - 120 cells each). Bars show mean ± SEM for each stimulation condition. Coefficient of determination (R²) and significance of linear regression shown for hippocampal subfields CA1, CA3 and dentate gyrus (DG) (****p<0.0001). **E)** Expression of cFos (arbitrary units) as a function of stimulation time. Circles represent averages from individual slice cultures, bars show mean ± SEM. Coefficient of determination (R²) and significance of linear regression is shown for each hippocampal subfield (CA1, ***p = 0.0009; CA3, ***p = 0.004; DG, **p = 0.0036). **F)** Correlations between CaMPARI2 photoconversion (Ca^2+^) and cFos expression for different stimulation durations. Mean ± SD of cells collected from all cultures (n = 3-9) per stimulation duration. (CA1 and CA3: ****p < 0.0001; DG: *p = 0.045). **G)** CaMPARI2 photoconversion (Ca^2+^) and cFos expression of individual neurons, color coded by stimulation time (same data as F). Note that for individual stimulation conditions, coefficients of determination are very low. **D-G)** For all correlations, Person’s correlation coefficient was computed. Similar results were observed in two independent experiments.

Sixty seconds of bicuculline stimulation with illumination resulted in H2B-CaMPARI2 photoconversion across all hippocampal subfields, indicating elevated nuclear Ca^2+^ levels during the stimulation period (Fig. 1B). This stimulation also led to widespread expression of cFos 60 min later (Fig. 1C), capturing the peak of endogenous cFos (Fig. S4). Quantitative analysis showed the strongest expression in CA1, DG and CA3, whereas neurons in CA2, hilus and subiculum expressed much less cFos (Fig. S5). To explore a range of activity levels, we varied the duration of bicuculline-driven activity by altering the timing of the TTX drop to terminate all electrical activity in the tissue. After fixation, CaMPARI2-positive nuclei were analyzed in a single image plane using Imaris (8 µm spots) to quantify CaMPARI2 conversion and cFos expression. Prolonging the period of high activity resulted in increased CaMPARI2 conversion (Fig. 1D) and increased cFos levels in DG, CA1 and CA3 (Fig. 1E). On the level of individual cultures, the correlation between CaMPARI2 conversion and cFos expression was very high (DG: R^2^ = 0.67; CA3 & CA1: R^2^ = 0.99, Fig. 1F). On the level of individual neurons, however, we found only weak or no correlations between these parameters at a given stimulation level (Fig. 1G). Even within a hippocampal subfield, neurons with intermediate photoconversion (0.4 to 0.6 [R/(R+G)]) showed a surprisingly large range of cFos levels, suggesting that there are additional parameters that control cFos expression in a cell-specific manner.

### cFos expression is dependent on the history of activity

Supraphysiological stimulation induced by cocaine consumption or epileptic seizures leads to cFos repression in the striatum and hippocampus (Cates et al. 2019; Renthal et al. 2008). Highly active regions share a common characteristic: they show substantial accumulation of ΔFosB, regardless of the specific stimulus. ΔFosB is a truncated variant of the FosB protein that has a very long lifespan. We evaluated ΔFosB expression levels at different time points after bicuculline stimulation. For each slice culture and hippocampal area, 20 neuronal nuclei were randomly selected based on the DAPI signal, their anti-ΔFosB fluorescence was evaluated and averaged (Fig. S6). Bicuculline stimulation induced strong ΔFosB expression in all hippocampal areas that lasted for at least 24 h (Fig. S6). To examine the interaction between ΔFosB and cFos, we stimulated cultures with a drop of bicuculline (10 µl, 20 µM) on day 1 (Fig. 2A). Since bicuculline is unstable at physiological pH and temperature, the period of increased activity is transient (Johnston 2013). The following day, the pre-stimulated cultures (referred to as ’BIC BIC’) were re-stimulated along with non-pre-stimulated cultures (’-BIC’). As in the previous experiments, activity was terminated by a drop of TTX after 60 seconds. After 60 min, the stimulated cultures were fixed and stained for cFos and ΔFosB. We analyzed the mean levels of cFos and ΔFosB expression across all neurons within each hippocampal subregion. The pre-stimulated cultures had reduced cFos levels in all subregions compared to the single stimulation condition (Fig. 2A, B). Conversely, ΔFosB levels were significantly higher in the pre-stimulated condition compared to the single-stimulated cultures (Fig. 2A, C). In double stimulated cultures (BIC BIC), all neurons had low cFos and high ΔFosB (Fig. 2D, black dots). Neurons stimulated once (-BIC) expressed cFos at higher levels (Fig. 2D, green dots). Similar results were obtained in CA3 (Fig. 2E-H) and CA1 (Fig. 2I-L). These results underscore that a period of highly synchronized activity can repress the cFos promoter in the majority of neurons 24 h later, coinciding with substantial ΔFosB accumulation. To establish a causal relationship between ΔFosB and cFos suppression at the single cell level, a more direct manipulation of ΔFosB was required.

**Figure 2:**
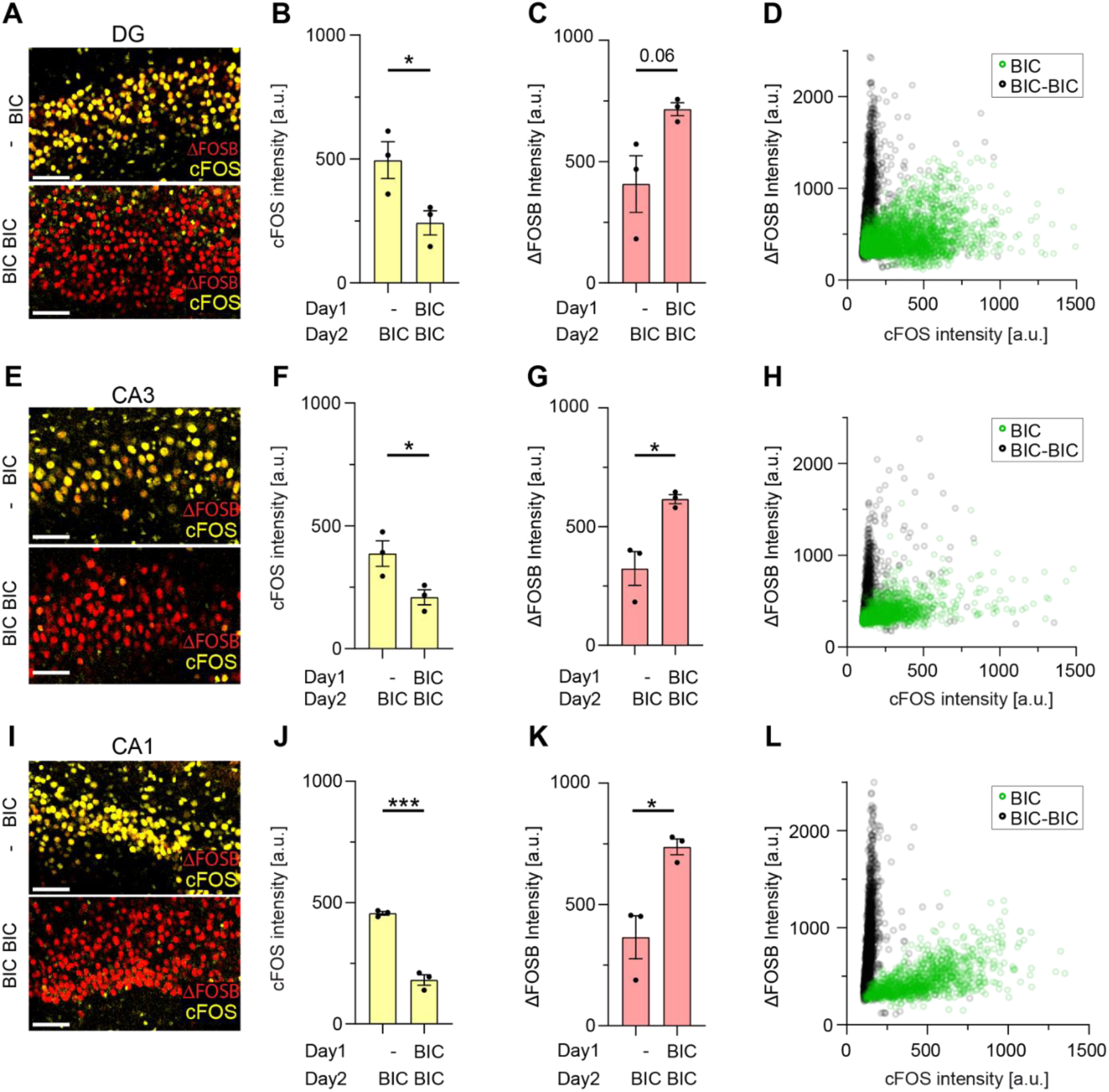
Bicuculline-induced activity increases ΔFosB expression in all hippocampal subregions. **A)** Confocal images of cFos (yellow) and ΔFosB (red) immunoreactivity in dentate gyrus (DG) after single (BIC) or two day-consecutive stimulation with bicuculline (BIC-BIC), scale bar 50 µm. **B)** Fluorescent intensity analysis (arbitrary units): cFos expression is decreased after consecutive (BIC-BIC) stimulation compared to single BIC stimulation (BIC) (*p = 0.046, t-test). Bars show mean ± SEM of all cells per culture (n = 3 slice cultures, black dots). **C)** Fluorescent intensity analysis (arbitrary units): ΔFosB expression is increased after consecutive (BIC-BIC) stimulation compared to single BIC stimulation (BIC) (ns p = 0.062, t-test). Bars show mean ± SEM of all cells per culture (n = 3 slice cultures, black spots). **D)** Single cell intensity plot of all cells of a single slice cultures per condition (green, BIC, n = 3177; black, BIC-BIC, n = 2605). **E-H)** Confocal images and analysis for CA3 neurons. cFos expression: *p = 0.042, t-test, BIC vs BIC-BIC. ΔFosB expression: *p = 0.016, t-test, BIC vs BIC-BIC. Single cell intensity plot: (green, BIC, n = 1265; black, BIC-BIC, n = 1219). **I-L)** Confocal images and analysis for CA1 neurons. cFos expression: ***p = 0.0003, t-test, BIC vs BIC-BIC. ΔFosB expression: *p = 0.017, t-test, BIC vs BIC-BIC. Single cell intensity plot: (green, BIC, n = 1077; black, BIC-BIC, n = 2081).

### Constitutive ΔFosB expression reduces activity-induced cFos in CA1

To generate ΔFosB overexpression in a few identified neurons, we electroporated individual CA1 neurons with ΔFosB (0.1 ng/µl) and cerulean (10 ng/µl), both under the control of the constitutive synapsin-1 promoter. Phosphorylation of ΔFosB at the serine 27 residue plays a critical role in extending its lifespan (Ulery-Reynolds et al. 2009; Ulery, Rudenko, and Nestler 2006). As a control, we therefore electroporated a modified version of ΔFosB in which the serine residue at position 27 was replaced with alanine (S-27A). After 6 days of expression, cultures were stimulated with bicuculline for 60 s, at which point all activity was terminated with a drop of TTX (Fig. 3A). Sixty minutes later, the tissue was fixed and stained for cFos, ΔFosB and DNA (DAPI) (Fig. 3B). Nuclei of electroporated neurons (DAPI + cerulean) and neighboring non-electroporated neurons (DAPI only) were selected independently of any cFos or ΔFosB signal (Fig. 3B). In a second step, the nuclear cFos and ΔFosB intensity of each electroporated neuron was normalized to the respective intensities of three neighboring neurons. This normalization procedure eliminated the influence of variable antibody staining between batches and variable activity levels between cultures. Both constructs (S-27 and S-27A) resulted in increased ΔFosB levels and decreased cFos expression compared to non-electroporated neurons (Fig. 3C). The wild-type variant (S27) resulted in significantly higher ΔFosB levels compared to the less stable S-27A version, leading to correspondingly lower cFos expression. The single cell intensity plot shows a strong inverse correlation between cFos and ΔFosB in S-27 expressing neurons (R = -0.59), which is not the case for the S-27A (R = - 0.04) or DAPI (R = 0.26) population. These results provide clear evidence for a causal relationship between ΔFosB expression and cFos repression in CA1 pyramidal cells. The single cell intensity plot (Fig. 3D) is reminiscent of the bimodal distribution observed in the populations of single or double bicuculline-stimulated cultures (Fig. 2L).

**Figure 3:**
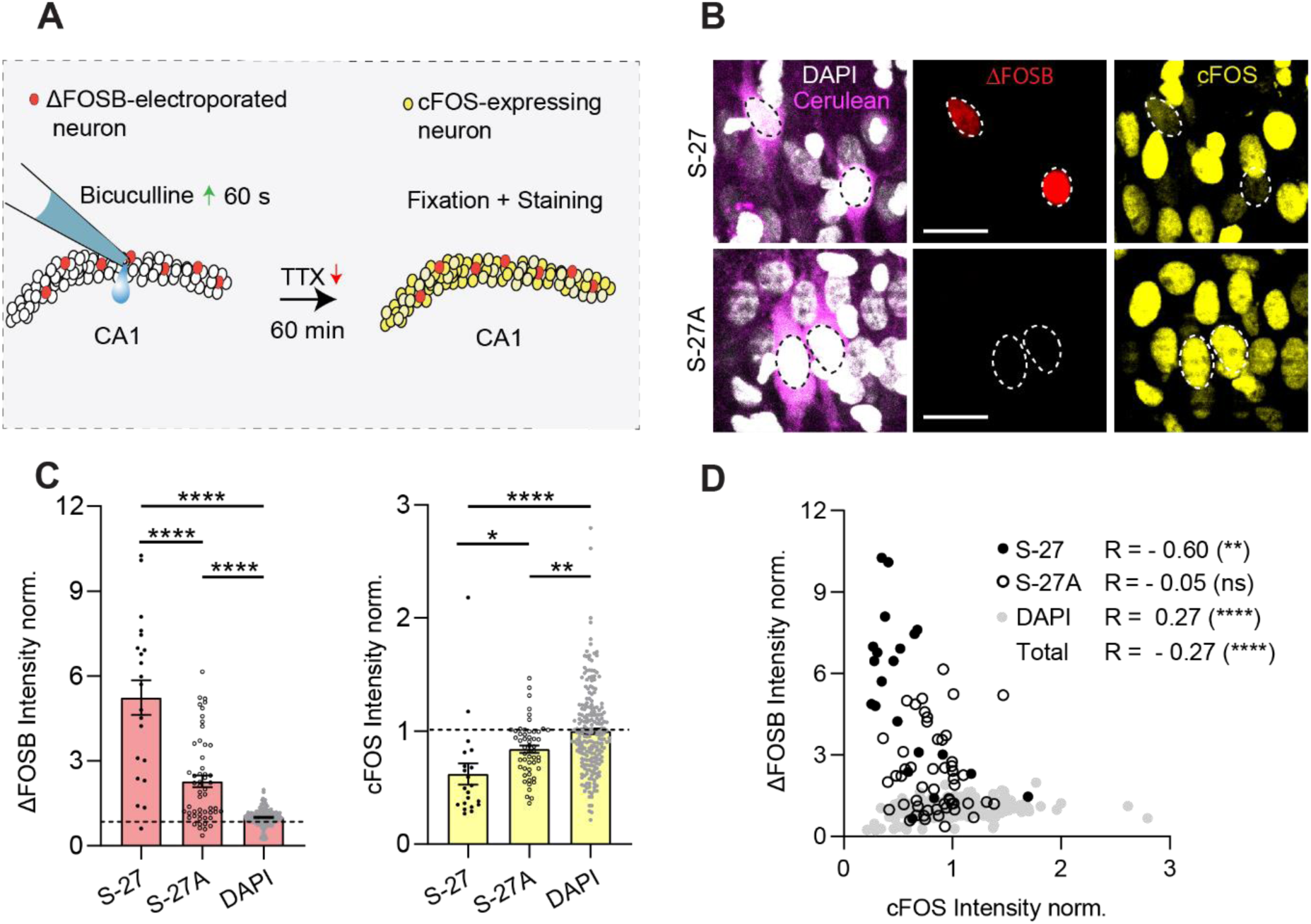
ΔFosB electroporation decreases bicuculline-induced cFos expression in CA1. **A)** Experimental design: slice cultures with single-cell electroporated CA1 neurons were stimulated with bicuculline for 60 seconds. Stimulation was terminated using TTX; cultures were fixed and stained for cFos (yellow) and ΔFosB (red) 60 minutes later. **B)** Confocal images of CA1 neurons expressing a non-modified version of ΔFosB (S27) and modified version of ΔFosB (S-27A) where the position serine 27 was exchanged with the amino acid alanine to prevent phosphorylation. In both conditions, neurons were co-electroporated with cerulean (10ng/µl, magenta) and the whole slice culture was stained for DNA (DAPI, gray) to allow unbiased cell selection for the intensity analysis. White circle show electroporated neurons. Scale bar = 20 µm **C)** Intensity analysis from neurons of n = 3 slice cultures per condition. Single spots represent a neuron and bars show mean ± SEM of all neurons. The cFos and ΔFosB intensities of electroporated neurons were normalized to neighboring DAPI^+^ neurons. Both protein versions (S-27, n = 21 neurons; S-27A, n = 53 neurons) show significantly higher ΔFosB expression (****p < 0.0001, S-27 vs DAPI; S-27A vs DAPI) compared to their neighboring DAPI^+^ neurons (n = 222). The wild type version leads to significantly more ΔFosB expression compared to the modified S-27A version (****p < 0.0001, S-27 vs S-27A). Neurons expressing both versions of ΔFosB have decreased cFos expression (****p < 0.0001, S-27 vs DAPI; **p = 0.01, S-27A vs DAPI) compared to neighboring DAPI^+^ neurons. Neurons expressing the S-27 version have decreased cFos intensities compared to the modified S-27A version (*p = 0.049, S-27 vs S27-A; 1-way ANOVA, Tukey’s multiple comparison). **D)** Single cell intensity plot and intensity correlations between cFos and ΔFosB of different conditions (Pearson’s). S-27 expressing neurons show a strong inverse correlation (R = - 60, ** p = 0.0043). S-27A (R = -0.05, ns p = 0.75), DAPI (R = 0.27, **** p < 0.0001), All (R = 0.27, **** p < 0.0001).

### ΔFosB represses cFos via HDAC recruitment

ΔFosB is a splicing product of FosB (Robison and Nestler 2022). When phosphorylated by casein kinase 2 (CK2), ΔFosB becomes resistant to degradation and accumulates for several days (Ulery-Reynolds et al. 2009; Ulery, Rudenko, and Nestler 2006). ΔFosB bineds directly to the cFos promoter and is able to recruit histone deacetylase class I enzymes (Renthal et al. 2008), setting the promoter to a repressed state (Fig 4A). We decided to test whether cFos suppression by bicuculline pre-stimulation (Fig. 2) indeed involves histone modification. Dentate gyrus neurons that were pre-stimulated with a drop of bicuculline (10 µl, 20 µM) 24 h before the second stimulation showed a CaMPARI2 conversion similar to that of non-prestimulated cultures, indicating that the network was still excitable (Fig. 4B, C). As in the previous experiments (Fig. 2), cFos expression was strongly suppressed by pre-stimulation (Fig. 4D). The HDAC inhibitor entinostat (MS 275) and the casein kinase II inhibitor DRB completely prevented the inhibitory effect of pre-stimulation in the dentate gyrus (Fig. 4D), but also increased nuclear calcium levels (Fig. 4C). Similar results were obtained in CA1 (Fig. 4E-J). In CA3, nuclear calcium levels were not affected by inhibition of CKII or HDAC (Fig. 4H,I), but cFos inhibition was largely prevented (Fig. 4J). Taken together, we demonstrated that epigenetic repression of cFos after high activity occurs in all hippocampal subfields and is likely mediated by phosphorylated ΔFosB.

**Figure 4:**
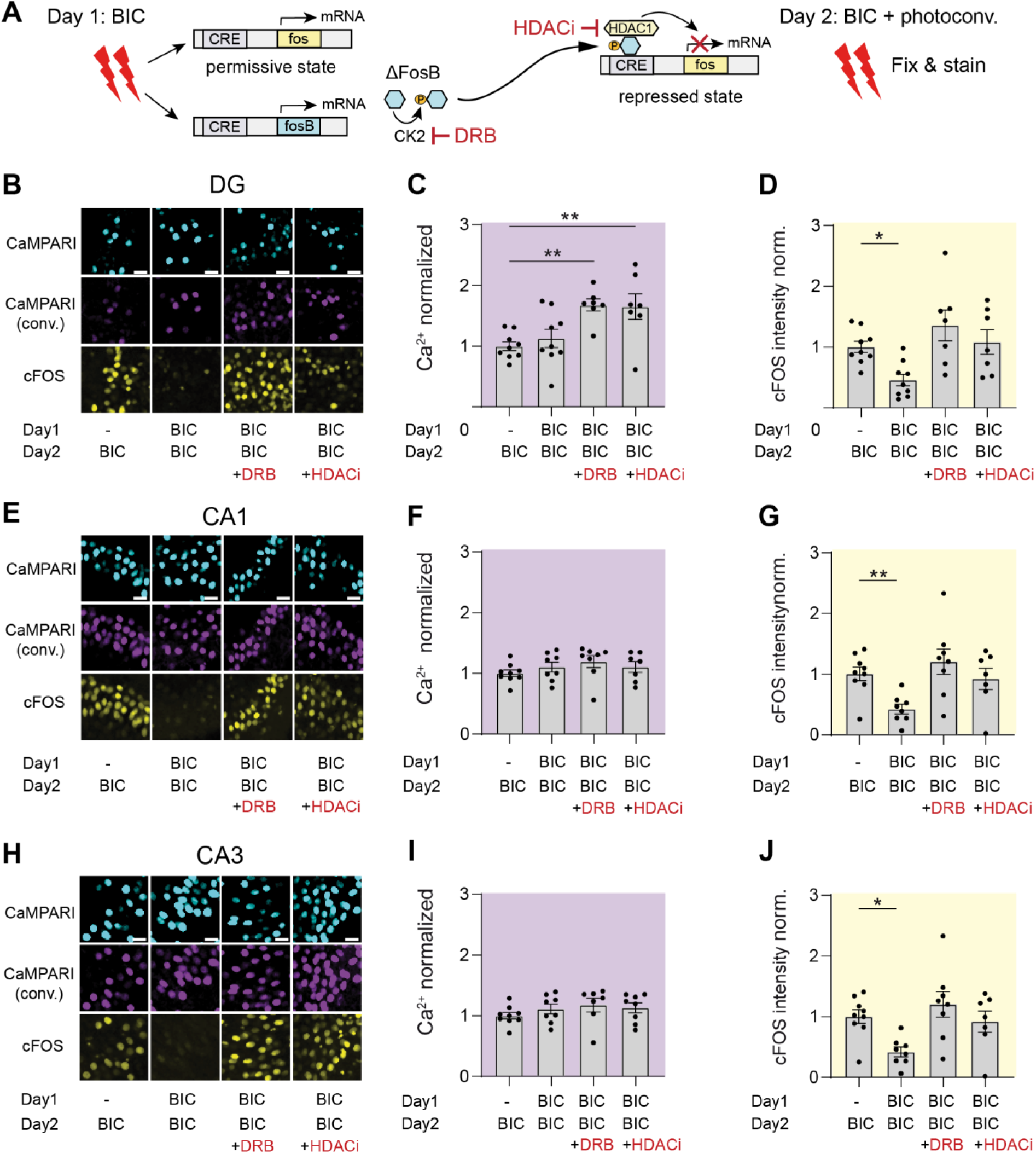
History of activity affects cFos expression. **A)** Bicuculline stimulation (BIC) on consecutive days, proposed mechanism of cFos inhibition. CK2, casein kinase 2; DRB, 5,6-Dichlor-benzimidazol-1-β-D-ribofuranosid; HDACi, histone deacetylase inhibitor Entinostat (MS 275). **B)** Confocal images of nuclear localized CaMPARI2 (native green fluorescence, cyan), photoconverted CaMPARI2 (immuno-enhanced, magenta) and anti-cFos staining (yellow) in dentate gyrus (DG), 1 h after bicuculline stimulation on day 2. Images show following conditions: BIC one-time stimulation on day 2 (BIC), BIC double stimulation on consecutive days (BIC-BIC), BIC double stimulation with CK2 blocker (DRB) added on day 1 (BIC-BIC + DRB), BIC double stimulation with HDACi (MS275) (BIC-BIC + HDACi) added on day 2 together with second bicuculline stimulation. **C)** Analysis of Ca^2+^ level in dentate gyrus (DG) based on CaMPARI2 photoconversion during 60 seconds of BIC stimulation on day 2. Black dots represent the mean intensity in arbitrary units of 20 randomly selected CaMPARI2^+^ nuclei. Gray bars represent mean ± SEM of n = 7 - 9 slice cultures per condition. All data points are normalized and statistically compared to the single stimulated condition (BIC). Double stimulated + HDACi or DRB treated cultures show increased calcium levels compared to BIC condition (**p = 0.005, BIC-BIC + DRB; **p = 0.0072, BIC-BIC + HDACi). Ca^2+^ level of double stimulated cultures were not different from the BIC condition (ns p = 0.84, BIC). **D)** cFos intensity analysis in arbitrary units. All data points are normalized and statistically compared to the single stimulated condition (BIC). BIC-BIC condition shows decreased cFos expression compared to the single BIC condition (*p = 0.044). BIC-BIC + DRB or BIC-BIC + HDACi show no significant difference in cFos expression compared to BIC condition (ns p = 0.30, BIC-BIC + DRB; *p = 0.97, BIC-BIC + HDACi). The regions CA1 and CA3 were analyzed the same way. **E)** Confocal images of CA1 region. **F)** Analysis of Ca^2+^ levels in CA1. All conditions show similar Ca^2+^ levels compared to BIC condition (ns p = 0.82, BIC-BIC; ns p = 0.16, BIC-BIC + DRB; ns p = 0.23, BIC-BIC + HDACi; n = 7 – 11 cultures). **G)** cFos intensity analysis in CA1. Only double stimulated condition shows decreased cFos expression compared to the BIC condition (**p = 0.0094, BIC-BIC; ns p = 0.88, BIC-BIC + DRB; *p > 0.99, BIC-BIC + HDACi) **H)** Confocal images of CA3 region **I)** Analysis of Ca^2+^ levels in CA3. All conditions show similar Ca^2+^ levels compared to BIC condition (ns p = 0.68, BIB-BIC; ns p = 0.22, BIC-BIC + DRB; ns p = 0.70 BIC-BIC + HDACi) **J)** cFos intensity analysis in CA3. Only double stimulated condition shows decreased cFos expression compared to the BIC condition (*p = 0.025, BIC-BIC; ns p = 0.66, BIC-BIC + DRB; ns p = 0.97 BIC-BIC + HDACi). Statistical test: 2-way ANOVA, Dunett’s multiple comparison.

### Physiological activity drives ΔFosB expression in the dentate gyrus, but not in CA1

The role of ΔFosB as a protein influencing the epigenetic machinery has been studied mainly in the context of pathological conditions or drug abuse. These conditions are associated with supraphysiological activity and its negative consequences, such as memory impairment and compulsive behavior. In a previous study, we evaluated cFos and ΔFosB expression in water maze-trained mice using a subtractive method to quantify ΔFosB expression (Lamothe-Molina et al. 2022). Here, we use a recently developed antibody that directly detects ΔFosB (Fig. 5A). After two consecutive days of water maze training, a similar number of neurons in DG and CA1 express cFos (Fig. 5B). However, the accumulation of ΔFosB is largely limited to granule cells in the DG (Fig. 5C). Single neuron analysis revealed two subsets of DG granule cells, displaying either high ΔFosB / low cFos or low ΔFosB / high cFos, with an inverse correlation between the two proteins (Fig. 5D). Only a small number of neurons expressing ΔFosB were detected in CA1, and this group also had low cFos levels (Fig. 5E). Therefore, although the inhibitory mechanism is present in CA1 pyramidal cells (Fig. 2), it appears to be minimally active during physiological activity.

**Figure 5:**
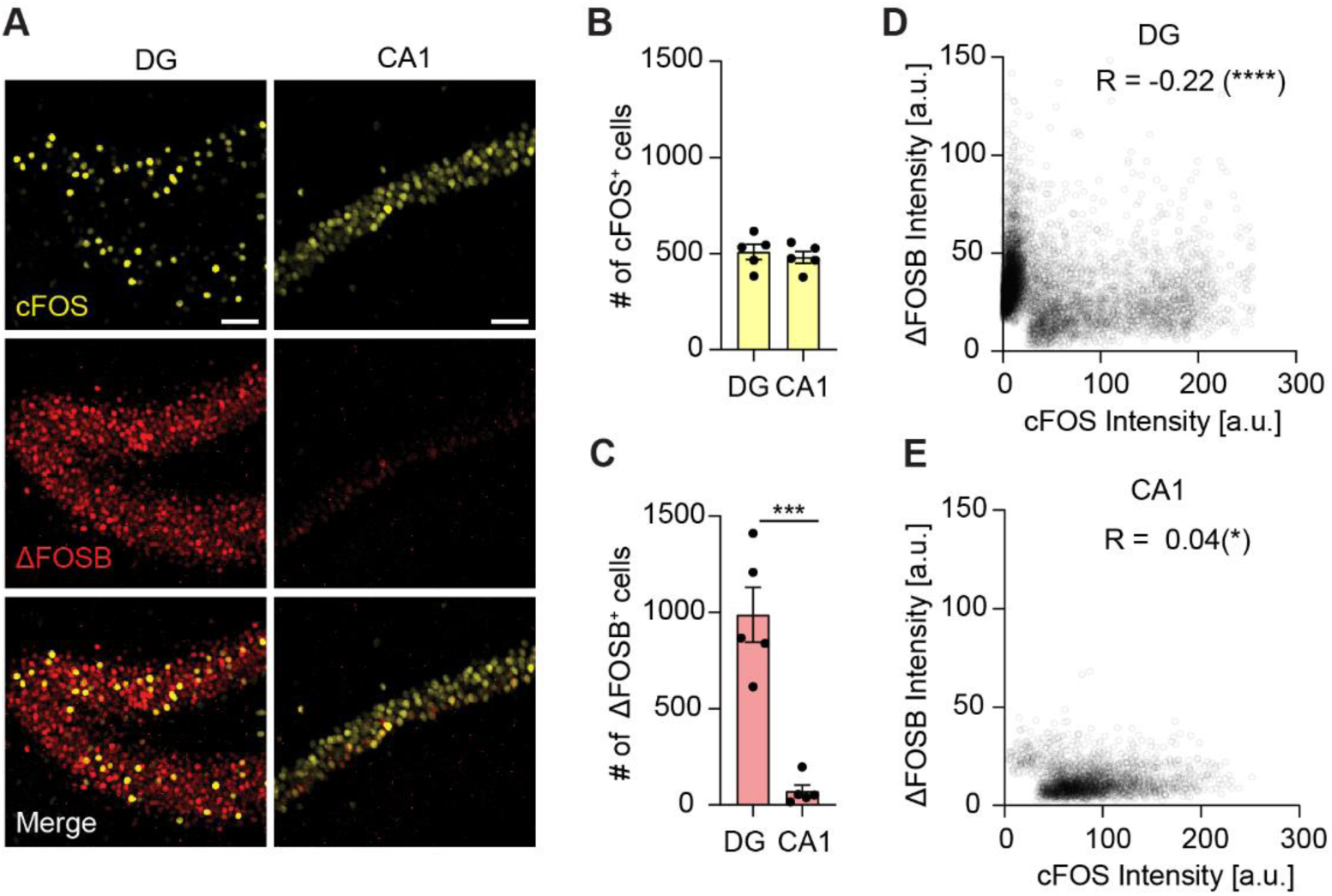
Expression of cFos and ΔFosB in the hippocampus after water maze training. **A)** In dentate gyrus (DG), sparse expression of cFos (yellow) and strong expression of ΔFosB (red). In CA1, dense expression of cFos and minor ΔFosB expression. Scale bar: 20 µm. **B)** No significant difference between cFos-expressing neurons in DG and CA1 (ns p = 0.60, t-test) after 2 days of water maze training (n = 5 mice). Dots show cell count (sum) of 4 analyzed images (18 µm z-stacks) per animal and region. Bars show mean ± SEM **C)** High number of ΔFosB-positive cells are found in DG, not in CA1 (***p = 0.0002, t-test) **D)** Single cell intensity analysis. Individual DG granule cell nuclei (circles) express either high cFos or high ΔFosB, rarely both (Pearson’s R = -0.22; **** p < 0.0001; n (cells) = 6617). **E)** CA1 pyramidal cells express cFos, but hardly any ΔFosB (Pearson’s R = 0.04; **** p < 0.0001; n (cells) = 2608).

### Effect of HDAC inhibition on cFos re-expression in the dentate gyrus

The strong expression of ΔFosB in the DG after water maze training and its repressive effect on cFos led us to ask whether we could counteract cFos repression by inhibiting histone deacetylase class I. For these experiments, we injected Tet-tag mice with AAV-TRE-mKate2 and kept them on doxycycline (Dox) to prevent cFos tagging during the recovery period (Fig. 6A). The day before the mice learned to locate a hidden platform in the water maze (day 1), doxycycline was removed from the food to permanently label cFos-expressing neurons with mKate2 (cFos tagging). After the training, further cFos tagging was prevented by an intraperitoneal injection of Dox. On day 2, mice were retrained to find the platform and perfusion-fixed 1 h later. To assess cFos expression on day 2, tissue was stained for cFos (2^nd^ cFos ensemble, 647 nm), and cFos-tagged GC from day 1 were enhanced with anti-mKate2 staining (cFos-tagged, 568 nm). To test whether HDAC inhibition affects cFos repression, one group of mice was injected with HDACi (Cl-994) immediately after training on day 1 and again 1 h before the training on day 2. The control group was injected with vehicle only (VEH, DMSO + 0.9% NaCl). As we and others previously reported, cFos expression in the DG was sparse on both days (Lamothe-Molina et al. 2022; Chatzi et al. 2019) (Fig 6B-C). HDACi treatment did not alter the size of cFos ensembles (Fig 5C, D). In HDACi-treated mice, 11% of day 1 Fos^+^ cells re-expressed cFos on day 2, compared to 7% in the control group (Fig. 5E). The above-chance re-expression of cFos in HDACi-treated animals was normalized to the control group (VEH) of each experimental round. The above-chance re-expression in HDACi-treated mice was significantly higher compared to the control group (Fig 6F), demonstrating that HDAC inhibition indeed affects cFos repression in dentate granule cells.

**Figure 6:**
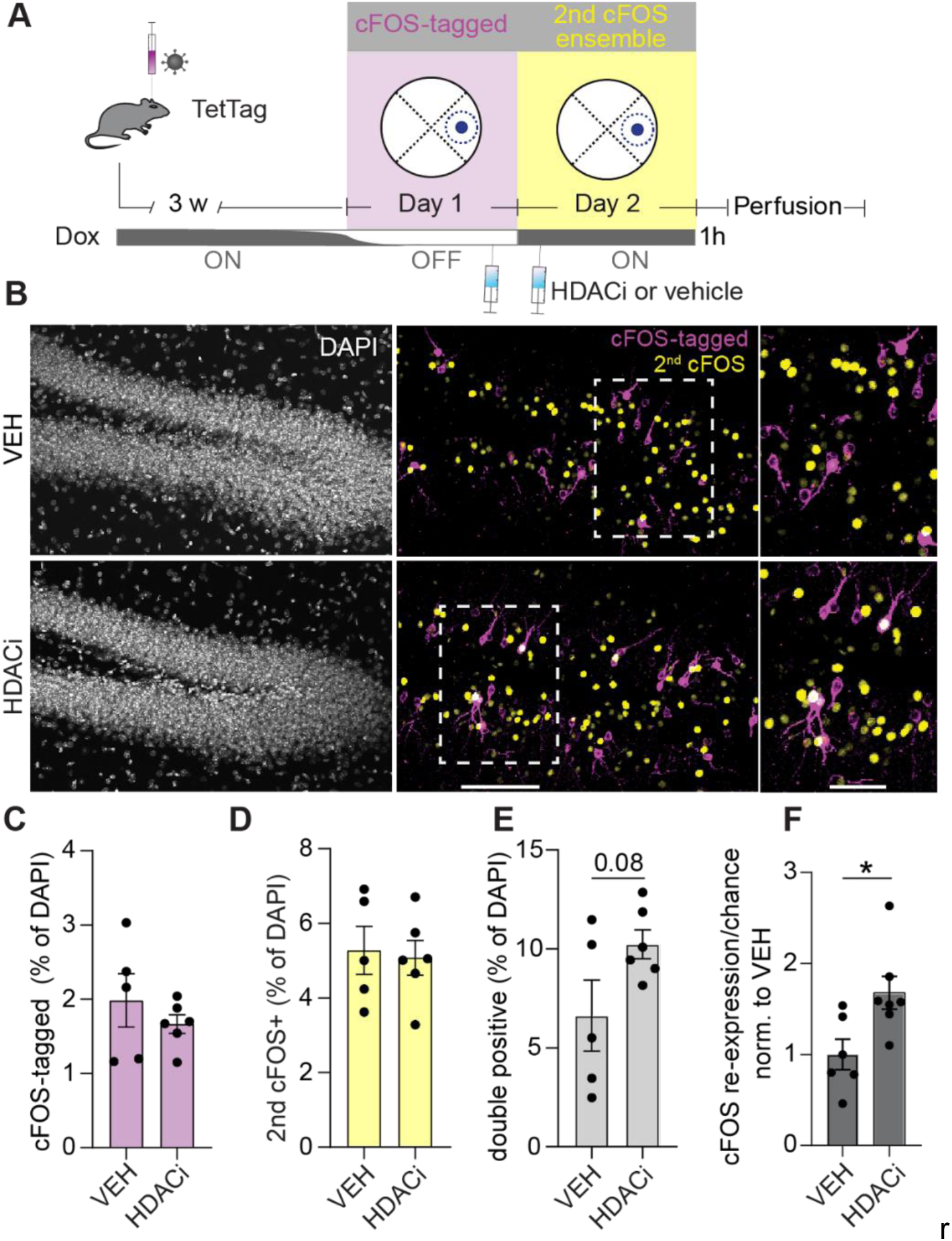
HDAC inhibitor increases cFos re-expression in dentate gyrus. **A)** Experimental time line: TetTag mice were injected with AAV-3TRE-mKate2 to tag cFos^+^ granule cells with mKate2 when doxycycline is removed from chow (OFF Dox). During the OFF Dox period (Day 1), mice were trained to find a platform in the water maze. On day 2, mice performed the same training protocol again (ON Dox) and were perfused 1 h after training. One hour after WM training on day 1 and 1 h beforeWM training on day 2, mice were injected with HDACi solution (Cl-994, 30 mg/kg in 90% 0.9% NaCl, 10% DMSO) or vehicle solution (90% 0.9% NaCl,10% DMSO). **B)** DAPI staining of dentate gyrus and immunofluorescence against mKate2 (magenta) and cFos (yellow). White square indicates zoomed section. **C)** No difference in percentage of granule cells expressing cFos on day 1 between HDACi and VEH treated animals (ns p = 0.37, t-test). **D)** No difference in percentage of granule cells expressing cFos on day 2 (ns p = 0.81, t-test). **E)** Increase in percentage of double positive GC in HDACi condition (ns p = 0.08, t-test). **F)** cFos re-expression above chance level normalized to VEH condition per experimental round, is significantly higher in HDACi-treated mice (*p = 0.019, t-test). 1^st^ experiment, n = 2 per condition; 2^nd^ experiment, n = 4 (HDACi), n = 3 (VEH).

### Inhibition of histone deacetylase during learning of multiple platform positions in the water maze

Multiple lines of evidence suggest that the DG plays a critical role in pattern separation, a key aspect of episodic memory and cognitive flexibility (Hainmueller and Bartos 2018; Leutgeb et al. 2007) Therefore, we tested whether the ΔFosB expression and cFos repression in the DG after water maze training is functional in facilitating reversal learning in the water maze, a cognitive ability known to rely on pattern separation and cognitive flexibility. To investigate the effects of HDAC inhibition during spatial reversal learning, we trained C57BL6/J mice to locate three different platform positions in the water maze over the course of 4 days. Some mice were orally treated with the HDAC inhibitor Cl-994 (HDACi-treated, n = 12 mice) one hour before water maze (WM) training on each training day, while others were administered the vehicle solution only (90% corn oil and 10% DMSO, n = 15 mice). All mice were trained for three days to find the platform in the east quadrant of the WM. The learning curve of HDACi-treated animals did not differ from that of control animals (Fig. 7A). In addition, both groups showed identical performance in the day 1 probe trial (Fig 7B), indicating no effect of HDACi on short-term memory. HDACi-treated mice performed again equally well in the probe trial the next day (Fig. 7C), suggesting that long-term memory was also unaffected. Further training improved performance, but revealed no effect of HDAC inhibition in probe trials (Fig. S7).

**Figure 7:**
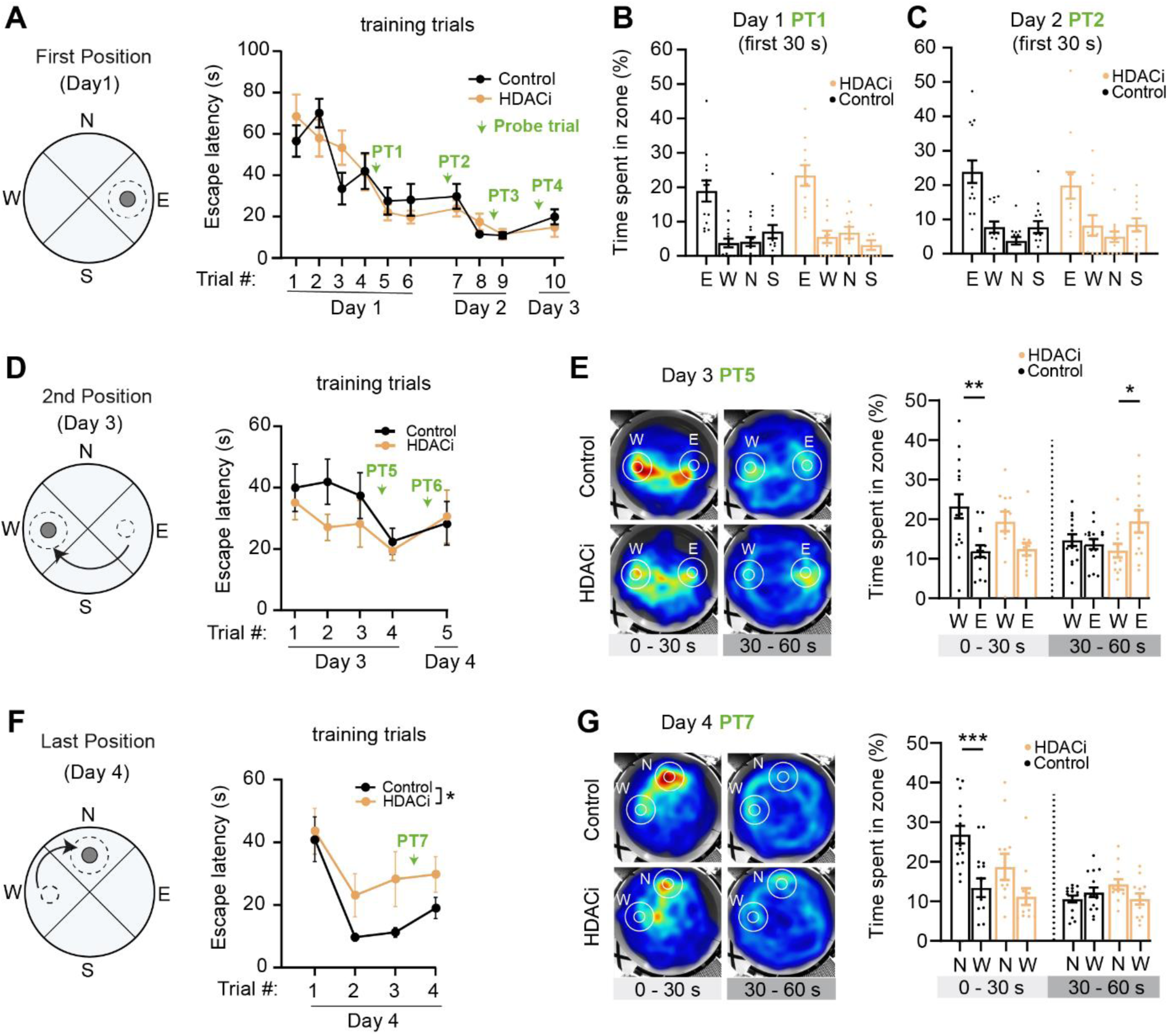
Spatial memory formation under HDAC inhibition. **A)** Day 1: The platform (gray circle) was located in the east quadrant of the maze. Escape latency to the platform was used for analysis of training trials (TT) and the dotted line (target zone, Ø = 42 cm) for probe trial analysis (PT). Mean escape latency (± SEM) improved over the course of 3 days during 10 TTs, with no difference between HDACi (ochre, n = 14-15) or VEH condition (black, n = 11-12) (ns, p = 0.99, 2-way ANOVA). Green arrows indicate probe trials. **B)** Performance during the first 30 s of the probe trail on day 1. Each point represents one animal (HDACi: n = 11, VEH: n = 13). Bars show percentage of time (mean ± SEM) spent in each target zone (east, E; west, W; north, N; south, S). No significant difference between HDACi and control group (ns p = 0.40, 2-way ANOVA). **C)** Probe trail performance (first 30 s) on day 2 to test for long-term memory. Bars show mean ± SEM. No significant difference between both conditions (ns p = 0.81, 2-way ANOVA). **D)** On day 3, mice were trained to a novel platform location (west) in four TTs. Escape latency to platform west is shown as mean ± SEM. No significant difference between HDACi and control group (ns p = 0.17, 2-way ANOVA) **E)** Mean swim trajectories of all animals per condition for the first (0 - 30 s) and second half (30 - 60 s) of the probe trial (heat maps). Animals alternate between the old (E) and new (W) target zones. Bars show mean ± SEM. During the first 30 s, VEH-treated animals (black) show a significant preference for the new learned location (W) compared to the first learned location (E) (**p = 0.0016, 2-way ANOVA, Šídák). HDACi-treated mice (ns p = 0.89, 2-way ANOVA). In the second half of the probe trial, HDACi-treated animals show a significant preference for the old (E) platform location (*p = 0.016, 2-way ANOVA). VEH-treated mice (ns p = 0.89, 2-way ANOVA). **F)** On day 4, mice were trained to yet another platform location (north) in four TTs. The VEH condition shows better learning performance (mean ± SEM escape latency) compared to the HDACi group (*p = 0.029, 2-way ANOVA). Green arrow indicates probe trial. **G)** Mean swim trajectories of all animals per condition for the first (0 - 30 s) and second half (30 - 60 s) of the probe trial (hear maps). Bars show mean ± SEM. Control animals show a significant preference for the new platform position (N) compared to the previous (W) position (***p = 0.0004, 2-way ANOVA, Šídák). HDACi-treated mice (ns p = 0.01, 2-way ANOVA). During 30 - 60 s no condition showed preference for platform positions north vs west (VEH, ns p = 0.51; HDACi, ns p = 0.09; 2-way ANOVA, Šídák).

Over the course of three days of training, mice developed a highly accurate memory for the platform location in the east quadrant. To test behavioral flexibility, mice underwent four reversal training trials to a new platform location on day 3 (platform in the west quadrant, Fig. 7D and E) and day 4 (platform in the north quadrant, Fig. 7F and G). After each reversal learning session, a probe trial was performed to evaluate the mice’s search strategies. The mice in the control group quickly adapted their search strategy to the new positions of the platform, as shown by their improved performance in the training trials and their behavior during the probe trials. By the fourth trial on day 3, the control group mice had improved their learning performance (Fig. 7D), and by day 4, they needed only one trial to locate the platform promptly at the new location (Fig. 7F). It is important to note that on Day 3, the mice had to learn two things in order to switch their search strategy: first, the new position of the platform, and second, the “rule” that the platform could only be found at the most recent location, not at the location where they had previously found it on most of the training trials. Therefore, it is possible that the mice’s rapid reversal learning on day 4 was facilitated by their prior understanding of this new rule. During the first 30 seconds of the probe trials, when mice are expected to search at the location with the higher probability of finding the platform, the control group spent more time at the newly learned platform location compared to the previously learned location (Fig. 7E and G, Fig. S8). Thus, the control mice exhibited adaptive behavioral flexibility and switched their strategy in response to the changing conditions experienced during reversal learning. In the HDACi-treated mice, however, this ability appeared to be partially impaired. They showed weaker performance during the reversal training on day 4 compared to the control mice (Fig. 7F), and they did not show a preference for the newly learned platform over the previously learned one during the probe trials on days 3 and 4 (Fig. 7E and G).

Taken together, these results suggest that HDAC inhibition does not affect the learning or recall of a single spatial location. However, when mice are presented with a choice between multiple possible platform locations, the spatial memory formed under HDAC inhibition (first location) appears to interfere with the storage or recall of new information, thereby reducing behavioral flexibility.

To further investigate the stability of spatial memories, all animals returned to the WM on day 5 for a long-term memory probe trial lasting 90 s. No HDACi was administered on this day. We split the total search path into three 30 s time periods and analyzed the number of entries into the three previously trained platform positions (Ø = 15 cm, Fig. 8A). In the first period, control animals had a strong preference for the last-learned platform position (green) compared to the second (red) and first (dark blue) position. Later in the probe trial (30 - 60 s), they sampled all three learned positions. In the last period (60 - 90 s), their search became more random, indicating that they adapted their strategy when they could not find the platform where they expected it to be. HDACi-treated mice (dotted lines) showed a slight preference for the last and second learned positions at the beginning of the trial (first period) and switched to random search in the later periods. Averaged over the total duration of the probe trial, HDACi-treated mice spent significantly less time in the 3 target zones (Ø = 42 cm) compared to controls (VEH vs HDACi, p = 0.02, Fig. 8B). In the first 30 s of the probe trial, control animals spend most of the time searching in the zone of the last trained platform position (Fig. 8C, D) while HDACi-treated mice did not. The effect of HDAC inhibition was even more pronounced when we tested the mice again on day 11, 6 days after the end of training (p = 0.001, Fig. 8E, F). The search strategy of control animals was remarkably consistent on day 5 and day 11 (Fig. 8G). After finding no platform at the last trained location, they went looking at the second and finally, at the first trained location. Mice that were trained under HDAC inhibition initially searched around the second trained location, but showed significant variance between day 5 and day 11 (Fig. 8H). In summary, the water maze experiments showed that HDAC inhibition during training of multiple platform positions leads to reduced performance in probe trials. Significant effects of HDACi treatment were detected immediately after training (Fig. 7) as well as 1, 4 (Fig S8) and 6 days later (Fig. 8).

**Figure 8:**
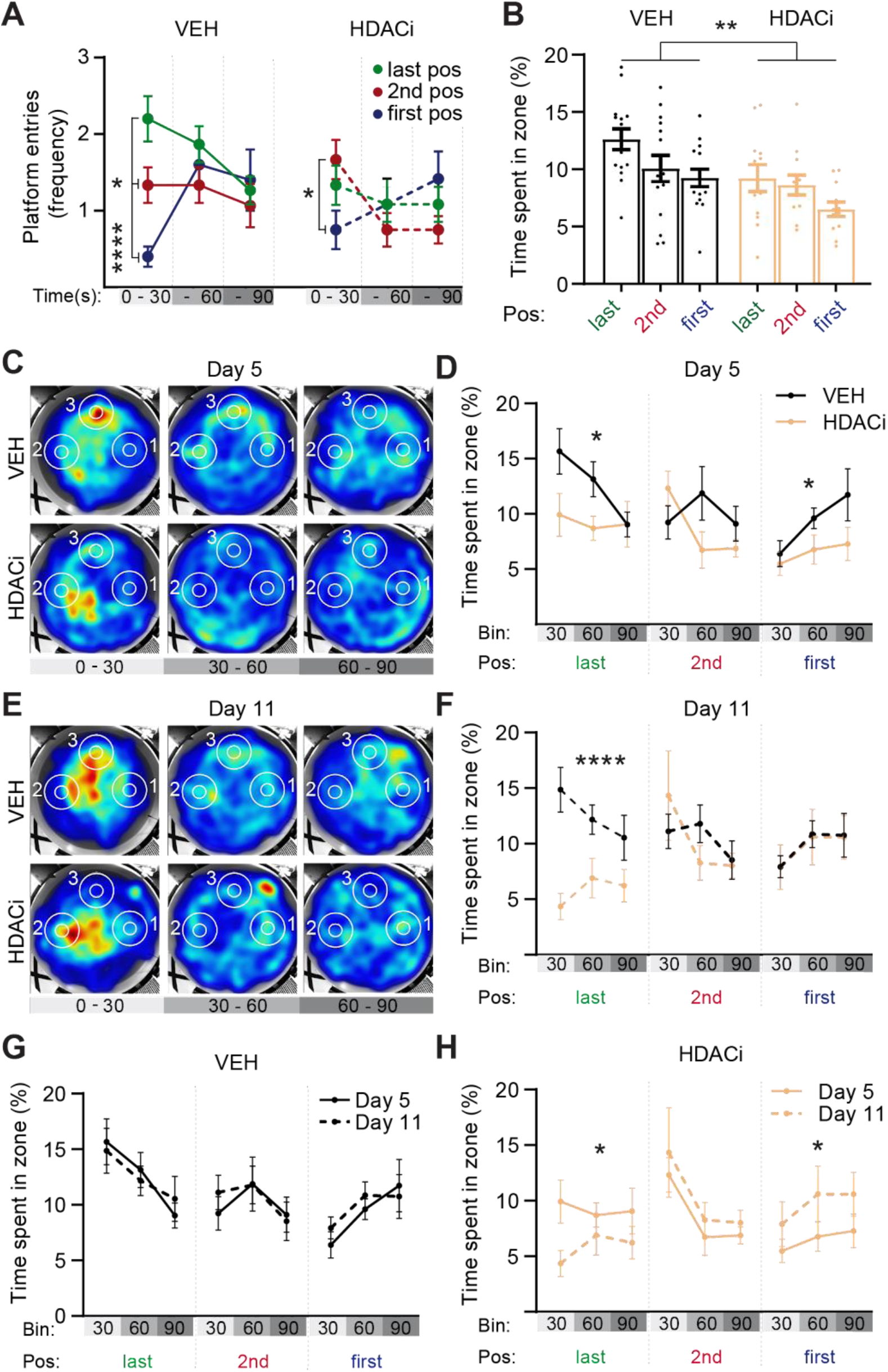
Spatial memory retrieval. **A)** Mean frequency of platform position entries for vehicle controls (VEH, n = 15 mice) and HDACi-treated mice (n = 12) during the probe trail on day 5. Last trained position (green, north), second trained position (red, west), first trained position (dark blue, east). The first 90 s were analyzed and divided into three periods: 0 - 30 s; 31 - 60 s; 61 - 90 s. Markers show mean ± SEM for each period. During the initial 30 s, control mice showed a strong preference for the last-trained platform location (last vs first, ****p < 0.0001; last vs 2^nd^, p = 0.051; 2^nd^ vs first, *p = 0.033, 2-way ANOVA, Tukey). Mice that were HDACi-treated during training showed less preference for a particular position (last vs first, p = 0.64; last vs 2^nd^, p = 0.26; 2^nd^ vs first, *p = 0.04, 2-way ANOVA, Tukey). **B)** Total percentage of time spent in target zone (last, 2^nd^, first). Each point represents one animal (VEH, n = 15; HDACi, n = 12), bars show mean ± SEM. HDACi-treated mice spent significantly less time in all target zones (VEH vs. HDACi, **p = 0.018, 2-way ANOVA). **C)** Mean trajectory projections (heat maps) of all animals on day 5, divided into three 30 s periods. **D)** Percentage of time spent in the three target zones per condition (mean ± SEM). Compared to the HDACi condition, VEH-treated animals show an increased preference for the last (north) and first (east) but not for the second (west) position (last, * p = 0.017; 2^nd^, p = 0.32; first, * p = 0.031; 2-way ANOVA, Šídák). **E)** Mean trajectory projections (heat map) of all animals per condition on day 11, divided in three 30-s periods. **F)** Percentage of time spent in the target zones (order: last, first, second) for all three periods per condition on day 11 (mean ± SEM). Compared to the HDACi condition, VEH-treated animals show a high preference for the last (north) but not for the second (west) or first (east) position (last, **** p < 0.0001; 2^nd^, ns p = 0.88; first, * p = 0.092; 2-way ANOVA, Šídák). **G-H)** Plots compare probe trail data of both days (5 vs. 11) per condition (VEH, HDACi). VEH treated animals show similar performance on both days (last, ns p = 0.95; 2^nd^, ns p = 0.78; first, ns p = 0.63; 2-way ANOVA, Šídák). Compared to day 5, on day 11, HDACi-treated animals spent less time around the last position (north, * p = 0.013) but more time around the first position (east, * p = 0.033), with no difference for the 2^nd^ learned position (west, ns p = 0.36; 2-way ANOVA, Šídák).

## Discussion

When we examined the relationship between nuclear calcium levels and cFos expression, we were struck by the poor correlation between these two parameters in individual neurons. The complex regulation of the cFos promoter has been studied in great detail (Robison and Nestler 2022; Vanhoutte et al. 1999), and we realized that accumulation of ΔFosB could have suppressed cFos specifically in neurons with a history of spontaneous activity. In hippocampal slice cultures, we were able to produce this effect by pre-stimulation, resulting in increased ΔFosB and decreased cFos in response to a second round of stimulation on the next day. Causality was established by observing the same inhibitory effect following overexpression of ΔFosB in single neurons. Long-lasting cFos inhibition required phosphorylation of ΔFosB by CK2 and histone deacetylation by HDAC1, as expected (Robison and Nestler 2022). In these slice culture experiments, we induced rhythmic epileptiform activity by sequential application of bicuculline and TTX, thereby transiently blocking GABA_A_-mediated inhibition. This supraphysiological stimulation caused ΔFosB accumulation and cFos inhibition in the DG, CA3, and CA1. Physiological activity, specifically water maze training, resulted in ΔFosB accumulation in DG granule cells, but not in CA3 and CA1 pyramidal cells. Thus, while the ΔFosB-based inhibitory mechanism appears to be operational and ubiquitous in all hippocampal neurons, spatial learning activates it specifically in the DG (Lamothe-Molina et al. 2022).

Our results indicate that ΔFosB accumulation in the DG during spatial learning restricts cFos expression to a subset of granule cells, namely cells that have not been strongly activated for several days. The specific activation of ΔFosB expression in the DG during spatial learning is intriguing given the role attributed to the DG in learning and memory. Multiple lines of evidence suggest that the DG plays a crucial role in pattern separation in both humans (Bakker et al. 2008; Berron et al. 2016) and rodents (Neunuebel and Knierim 2014; Madar, Ewell, and Jones 2019). Pattern separation allows for the discrimination between similar stimuli and memories, which is essential for cognitive flexibility and the ability to shift strategies when conditions change. A standard measure of cognitive flexibility and strategy shifting is reversal learning. We speculated that learning-induced accumulation of FosB in granule cells may be relevant for reversal learning in the water maze.

To test this hypothesis, we developed a memory task that required memorization of three different spatial locations, each of which was trained on a different day. Since histone deacetylation was necessary to produce ΔFosB-mediated cFos inhibition in culture, we used an HDAC1 inhibitor to disable this epigenetic mechanism in vivo. Although both groups of mice were able to learn and remember a single platform position, HDAC-inhibited mice had significant difficulty remembering novel positions (reversal learning). Previous studies have reported improvements in memory following HDAC inhibition (Peixoto and Abel 2013; Ramirez-Mejia et al. 2021; Villain, Florian, and Roullet 2016) The deficit we report here is specific to reversal learning and can thus be reconciled with the aforementioned studies. Obviously, increased memory persistence could lead to reduced behavioral flexibility when an animal is faced with more complex tasks and changing environments. Furthermore, the duration of experimental HDAC inhibition appears to be critical, as acute and chronic inhibition have different effects on addiction memory (Nestler 2014).

In our experiments with TetTag mice, HDAC inhibition increased the proportion of granule cells expressing cFos on two consecutive days of water maze training. In these experiments, we used a relatively simple task of training TetTag mice to a single platform position. It remains to be seen how more complex tasks affect the overlap of cFos populations in the dentate gyrus with and without HDAC inhibition. The effects of HDAC inhibition on memory are due changes in the expression level of genes regulated by the CREB:CBP transcriptional complex (Vecsey et al. 2007). Despite this relatively specific effect, it should be noted that ΔFosB-mediated inhibition of cFos is not the only memory-relevant process controlled by HDAC1. Calbindin-D28k has also been shown to be downregulated by ΔFosB-mediated histone deacetylation (You et al. 2017).

The behavior of reversal-trained mice was intriguing and changed during the probe trials. Control mice not only learned the new platform locations as indicated by the reversal learning trials on days 3 and 4), but also changed their strategy. During the first reversal learning on day 3, mice still seemed to follow the rule “search at the position where you found the platform most often”, as their performance slowly improved over the four training trials. In contrast, on day 4, when they underwent a second set of reversal training trials, control mice started swimming directly to the new platform position after only one training trial, indicating a new behavioral rule: “search at the position where you last found the platform”. In addition, our analysis of probe trials on days 5 and 11 suggests that the control group of mice remembered platform locations in a sequential manner (Fig. 8G). An anthropomorphic interpretation of this ordered search could be a cognitive process that takes into account new information, such as the absence of the platform at the expected location: “Given that the platform is not where I last found it, where else could it be?” The reuse of the same set of neurons for memory storage on consecutive days could hinder the ability to recall different locations in sequence while updating prior beliefs about the platform’s location. This is evidenced by the behavior of mice trained under HDAC inhibition, which did not visit the platform locations in reverse order (Fig. 8H), suggesting a lack of clearly structured memory regarding possible escape locations.

ChIP-sequencing of mouse hippocampus revealed that ΔFosB regulates genes involved in neuron and dendrite development and synapse maturation (You et al. 2018), perhaps in a homeostatic manner to prevent seizure generation (Eagle et al. 2018). Our results highlight the importance of epigenetic mechanisms in the organization of complex contextual memories (Creighton et al. 2020; Eagle et al. 2015; Ramamoorthi et al. 2011; Weng et al. 2018) and explain why calcium levels are not sufficient to predict cFos activation in individual neurons. In contrast to CA1 pyramidal cells, DG granule cells exhibit a dramatic shift in transcription upon activation. The common perception of the DG as a quiet, sparsely active region is accurate in describing its electrophysiological responses, but not in describing its transcriptional responses (Jaeger et al. 2018). While there is strong electrical reactivation of granule cells when exposed to the same environment on consecutive days, readily measurable by monitoring somatic [Ca^2+^]_i_ (Hainmueller and Bartos 2018), the state of intracellular signaling pathways is completely altered compared to the first exposure (Coba et al. 2012; Jaeger et al. 2018). Granule cells are the only neurons in cortex that require regular replacement, and it is tempting to speculate that their high transcriptional activity and associated increases in excitability wear them down. The dentate gyrus evolved rapidly during the transition from reptiles to stem mammals (Hevner 2016), undoubtedly to support novel memory or thought processes specific to mammals. Determining how the epigenetic state of granule cells affects their input-output function is the next challenge in understanding the unique computations that require such unusual hardware.

## Methods

### Organotypic hippocampal slice cultures

Wistar rat pups were sacrificed at postnatal day 5-7, both hippocampi were dissected and cut into 400 µm thick sections using a tissue chopper. The hippocampal slices were cultured on 30 mm sterile membrane inserts (Millipore) in 6-well plates, each containing 1 ml of culture medium, in a cell culture incubator (37°C, 5% CO_2_). The culture medium, consisting of 394 ml Minimal Essential Medium (Sigma), 100 ml heat-inactivated donor horse serum (Sigma), 1 mM L-glutamine (Gibco), 0.01 mg ml-1 insulin (Sigma), 1.45 ml 5 M NaCl (Sigma), 2 mM MgSO_4_ (Fluka), 1.44 mM CaCl_2_ (Fluka), 0.00125% ascorbic acid (Fluka), 13 mM D-glucose (Fluka), was partially (∼70%) replaced twice a week. Antibiotics, which can induce seizure-like activity in hippocampal slice cultures, were not required due to sterile preparation conditions.

### Single-cell electroporation

At DIV 14-16, CA1 neurons from slice cultures were transfected via single-cell electroporation (Wiegert, Gee, and Oertner 2017). Thin-walled pipettes (∼10 MΩ) were loaded with plasmids diluted in intracellular solution (Table 1). Under visual control (IR-DIC), the pipette was advanced against the neuronal membrane. DNA was ejected with 12 hyperpolarizing pulses (−12 V, 0.5 ms) using an Axoporator 800A (Molecular Devices). Experiments were conducted 6 days after electroporation.

**Table 1:** Plasmid list

| Plasmid | Short name | Concentration<br>(final) |
| --- | --- | --- |
| pAAV-hsyn1-H2B-CaMPARI2 | H2B-CaMPARI2 | Virus production |
| pAAV-CMV-ΔFosB<br>(Addgene plasmid 68545) | ΔFosB(S27) | 0.1ng/μl |
| pAAV-CMV-ΔFosB(S27A) | ΔFosB(S27A) | 0.1ng/μl |
| pCI-syn1-CFP |  | 10ng/μl |

### Viral vector production, dilution and transduction

Recombinant AAV were produced in the vector core facility of the University Medical Center Hamburg-Eppendorf. Virus particles were diluted to a working concentration of 1.13 x 10^12^ vector genomes / ml (vg/ml) in buffer solution containing (in mM) 145 NaCl, 10 HEPES, 25 D-glucose, 1.25 NaH_2_PO_4_, 2.5 KCl, 1 MgCl_2_, 2 CaCl_2_. In a laminar flow hood, 2 µl of the virus suspension was dropped on top of each slice culture at day in vitro (DIV) 14-19. After 5 days of expression, slice cultures were used for experiments.

### Bicuculline-induced stimulation and H2B-CaMAPRI2 photoconversion in the incubator

Slice cultures were either virally transduced or electroporated with plasmid DNA before stimulation. Bicuculline (BIC) was applied to induce activity to analyze Ca^2+^ levels (H2B-CaMPARI2), cFos and ΔFosB expression (see Table 2). H2B-CaMPARI2 expressing cultures were illuminated with 395 nm light (0.4 mW/mm^2^) during BIC stimulation. Three hippocampal slices were placed near the center of a single culture membrane in a 35 mm petri dish and transferred to the stimulation incubator at day 4 after transduction. At day 5 after transduction, the culture dish was placed into an LED illumination tower inside the incubator. To induce activity, slice cultures were treated by pipetting 100 µl bicuculline (BIC, 20 µM) on top of the cultures. After 10 - 60 s, tetrodotoxin (TTX, 300 µl, 1 µM) was added to prevent further spiking. The 395 nm LED (0.4 W/mm^2^) was turned on 5 s before BIC application and turned off 5 s after TTX application to ensure conversion during potential activity bursts induced by handling. After 60 min of incubation, slice cultures were fixed and stained (see IHC section).

**Table 2:** Slice culture experiments

| Figure | Virus/plasmid | BIC Stimulation |  | Stim. time | AB staining |
| --- | --- | --- | --- | --- | --- |
|  |  | Day 1 | Day 2 |  |  |
| 1 | H2B-CaMPARI2 | No | Yes | 10-60 s | cFos, CaMPARI2 |
| 2 | none | No | Yes | 60 s | cFos, ΔFosB |
| 3 | S27 / S27A | No | Yes | 60 s | cFos, ΔFosB |
| 4 | H2B-CaMPARI2 | Yes | Yes | 60 s | cFos, CaMPARI2 |

### Bicuculline stimulation on two consecutive days, DRB (CK2i) and MS-275 (HDACi) treatment

Slice cultures expressing H2B-CaMPARI2 were pre-stimulated with a drop of bicuculline (BIC, 10 µl, 20 µM, 2.7% DMSO, 97.3% buffer) the day (day 4) before the photo-conversion in the stimulation incubator. Additionally, some slice cultures were treated with BIC and 5,6-Dichlor-benzimidazol-1-β-D-ribofuranosid (DRB, Producer) (10 µl, BIC, 20 µM, DRB, 120 µM, 2.7% DMSO, 97.3% buffer). On day 5 slice cultures were stimulated with BIC again (100 µl, 20 µM, 2.7% DMSO, 97.3% buffer) in the presence of continuous 395 nm (0.4 mW/mm^2^) light. Some conditions were treated with BIC and MS-275 (HDACi, Producer) (100 µl, BIC, 20 µM, MS-275, 10 µM, 2.7% DMSO, 97.3% buffer). 60 s after BIC application, TTX (300 µl, 1 µM) was applied to abolish activity and the cultures were fixed and stained 1 hour later (see IHC section).

### Tissue fixation and Immunohistochemistry (IHC)

Slice cultures were fixed in 4% PFA/PBS solution for 25 min. Fixed cultures were cut off the membrane and washed in 1 × PBS for 10 min, incubated in blocking solution (500 µl, 1 × PBS, 0,3 % TritonX (Sigma-Aldrich), 5 % goat serum) for 2 h, then incubated in the primary antibody (Table 3) carrier solution (500 µl, 1 × PBS, 0,3 % Triton X-100, 1 % goat serum, 1 % BSA) overnight at 4°C. The next day, slice cultures were washed three times with 1 × PBS for 5 min, then incubated for 2 h in the secondary antibody (Table 3) carrier solution (500 µl, 1 × PBS, 0,3 % Triton X-100, 1 % goat serum, 1 % BSA) at RT. Slice cultures were washed 3 × 10 min in 1 × PBS, optionally stained with 4′,6-diamidino-2-phenylindole (DAPI, 1:1000) for 5 min, and mounted on coverslips using Immu-Mount (Shandon). This protocol was also used to stain acute hippocampal slices.

**Table 3:**
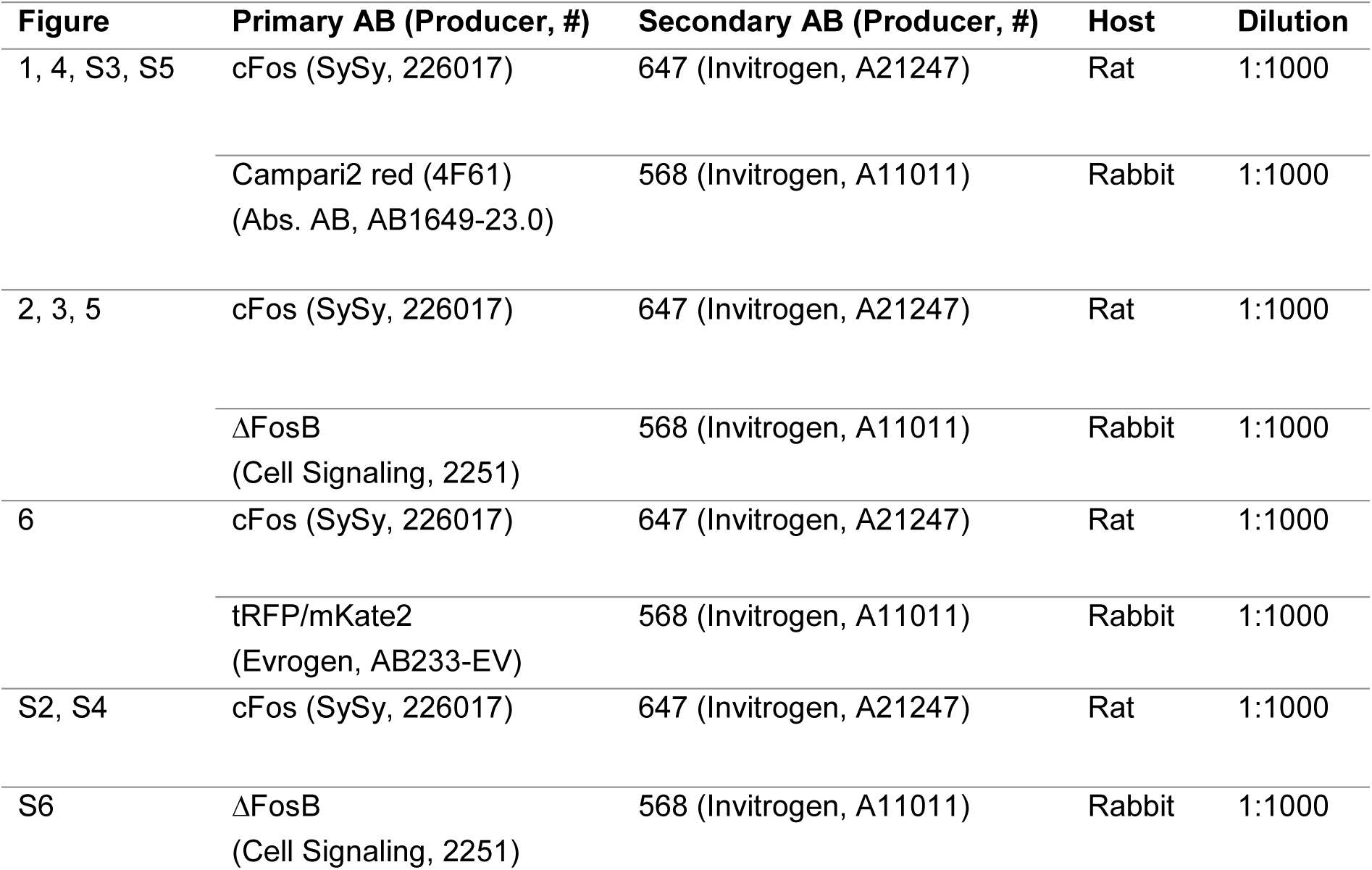
Primary and secondary antibody combinations (SySy, Synaptic Systems)

### Confocal imaging

To evaluate all hippocampal subfields (Fig. 1B-C, Fig. 2, Fig. S5), immunostained slice cultures were imaged in 3D with a confocal microscope (Olympus Fluoview FV 1000, UPLSAPO 20x/0.85) in 5×4 mosaic mode. Each stack contained up to 100 images (512 × 512 pixels, z-step: 3 µm). The following filter sets were used: Alexa 488 (CaMPARI2), Alexa 568 (CaMPARI2-converted, ΔFosB), Alexa 647 (cFos). Image stacks were stitched with ImageJ (Fig. 1A, B) (Preibisch, Saalfeld, and Tomancak 2009). Single plane images of hippocampal subregions (Fig. 1D-H, Fig. 4, Fig. S1, Fig. S2, Fig. S3, Fig. S4, Fig. S6) were acquired with a Zeiss LSM 900 (Plan-Apochromat 20×/0.8, 512 × 512 pixels). To image ΔFosB-electroporated CA1 neurons (Fig. 3), stacks of 7-9 image planes (1024 × 1024, z-step: 3 µm) were acquired using filter sets for Alexa 405 (DAPI, Cerulean), Alexa 488 (CaMPARI2), Alexa 568 (CaMPARI2-converted, ΔFosB), and Alexa 647 (cFos). The image acquisition settings were kept constant for all images within an experimental data set.

### Imaris spot selection and analysis

For automated or manual selection of nuclei in single plane or image stacks, we used the Imaris Spot Detection tool (see Table 4). For automatic spot selection, the quality threshold was kept constant for all images within an experimental data set. Spots detected outside the principal cell layer (corresponding to inhibitory neurons) were manually removed. For single plane zoomed-in images, 20 spots were manually selected based on the green CaMPARI2 signal or the DAPI signal (depending on the experiment, see Table 4). The color channels containing the measured intensities (cFos, CaMPARI2 converted, or ΔFosB) were turned off to avoid biased spot selection. For background subtraction in CaMPARI2 experiments, 10 additional spots were manually placed in background regions (all color channels turned on). The mean intensity of background spots (green and red channel) was subtracted from the spot intensities of non-converted (green) and converted (red) CaMPARI2 nuclei using a Matlab script. For the experiment shown in figure 3, all electroporated CA1 neurons were selected based on the cerulean signal. For each electroporated neuron, three neighboring non-electroporated neurons were manually selected based on the DAPI signal. The nuclear intensity (cFos, ΔFosB) of electroporated CA1 neurons was normalized to the mean nuclear intensity of the neighboring DAPI^+^ neurons (Matlab) to correct for variability in staining and bicuculline-induced stimulation intensity per culture.

**Table 4:**
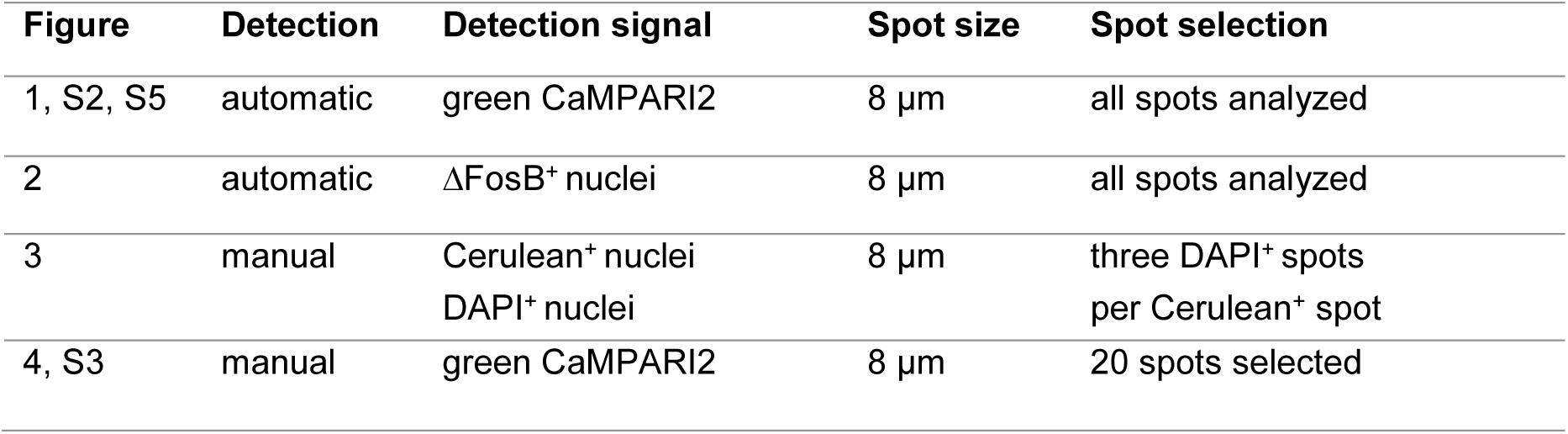
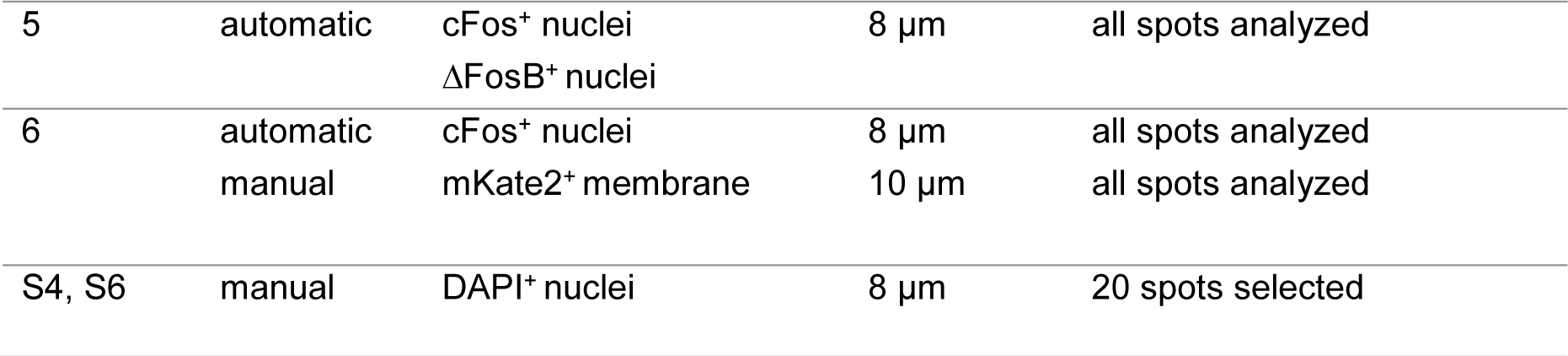
Imaris spot selection methods

| Figure | Detection | Detection signal | Spot size | Spot selection |
| --- | --- | --- | --- | --- |
| 1, S2, S5 | automatic | green CaMPARI2 | 8 μm | all spots analyzed |
| 2 | automatic | ΔFosB<sup>+</sup> nuclei | 8 μm | all spots analyzed |
| 3 | manual | Cerulean<sup>+</sup> nuclei<br>DAPI<sup>+</sup> nuclei | 8 μm | three DAPI<sup>+</sup> spots<br>per Cerulean<sup>+</sup> spot |
| 4, S3 | manual | green CaMPARI2 | 8 μm | 20 spots selected |
| 5 | automatic | cFos<sup>+</sup> nuclei<br>ΔFosB<sup>+</sup> nuclei | 8 μm | all spots analyzed |
| 6 | automatic<br>manual | cFos<sup>+</sup> nuclei<br>mKate2<sup>+</sup> membrane | 8 μm<br>10 μm | all spots analyzed<br>all spots analyzed |
| S4, S6 | manual | DAPI<sup>+</sup> nuclei | 8 μm | 20 spots selected |

Spot intensities were exported in csv files and further analyzed in Matlab. Nuclear [Ca^2+^] was estimated by calculating CaMPARI2 conversion after fixation and immunostaining as

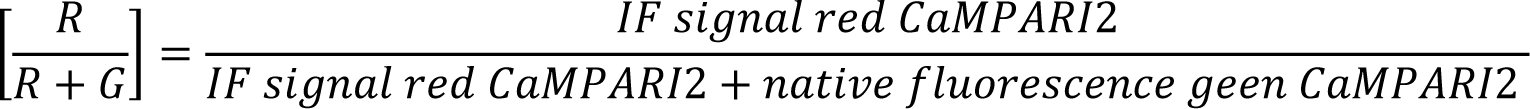

yielding values between 0 (no conversion) and 1 (complete conversion).

### Experimental animals

B6.Cg-Tg^(Fos-tTA,Fos-EGFP*)1Mmay/J^ (TetTag) mice were obtained from the Jackson Laboratory (Strain #018306) and bred to wild type (non-carrier) C57BL6/J mice from our colony. C57BL6/J (Jackson stock #:000664) were obtained from the UKE animal facility. Mice were group-housed with littermates until 2 weeks before rAAV injection or WM training and were then single caged. Mice had access to food and water ad libitum and were kept in an animal facility next to the behavioral rooms on a reversed light-dark cycle (dark 7 am - 7 pm) at 20 -23 °C with 45-65% humidity. Behavioral experiments were conducted during the dark phase of the cycle. Male and female TetTag mice between 20 - 40 weeks were included in the ΔFosB expression analysis (Fig 5) and cFos re-expression experiments (Fig 6). Male C57BL6/J mice between 20-40 weeks were used for HDAC inhibition experiments (Figs. 7 and 8). All experiments were conducted in accordance with German law and European Union directives on the protection of animals used for scientific purposes and were approved by the local authorities of the city of Hamburg (Behörde für Justiz und Verbraucherschutz, Lebensmittelsicherheit und Veterinärwesen, N 100/15 and N 046/2021).

### Stereotactic virus injection

TetTag mice were virus-injected under analgesia and anesthesia using a stereotaxic drill and injection robot (Neurostar). Mice were fixed to the frame under isoflurane anesthesia (1.5% mixed in O^2^), skin and connective tissue was removed, and two craniotomies were performed using an automated drill at the desired coordinates. We injected AAV_PHP.eB_ (10^12^ vg/ml) encoding a red fluorescent protein fused to an opsin (pAAV-TRE3G-BiPOLES-mKate2 (Addgene # 192579)) to label the membrane of cFos^+^ neurons. For bilateral injections into the dorsal hippocampus, we used a glass micropipette attached to a 5 μl syringe (Hamilton). A single injection per site was performed using stereotaxic coordinates for DG (−2.2 AP, ± 1.37ML, −1.9 DV) with a volume of 500 nl on each side (injection speed: 100 nl/min). After the last injection, the pipette was retracted 200 µm and left for at least 5 min to minimize efflux of virus during withdrawal. After the injections, the bone surface was cleaned with 0.9% NaCl solution and the skin was stitched. To avoid hypothermia, a heated pad was placed under the animal during surgery and under its cage for 1 h until full recovery. We provided post-surgery analgesia with Meloxicam mixed with softened Dox-food (see below) for 3 days after surgery. Animals recovered at least 2 weeks before behavioral experiments.

### Doxycycline treatment

Animals were given doxycycline-containing food (Altromin-Dox, 50 mg per kg of body weight, red pellets). To tag the first cFos ensemble, animals were changed to doxycycline-free food (Altromin, light-brown pellets) 24 h before exposure to the water maze. Doxycycline (50 µg/g bodyweight) was injected I.P. immediately after WM training on day 2.

### HDACi injection and oral delivery

For cFos re-expression analysis, Tet-Tag mice were I.P. injected with HDACi solution containing Cl-994 (30 mg/kg, Tocris #2952), 30% Kolliphor (Sigma-Aldrich) and 10% DMSO (Roth) in 0.9% NaCl solution. The first injection was given on day 1 immediately after WM training (OFF-Dox) and again one hour before the first training trial on day 2. The control group was injected with vehicle solution (10% DMSO, 30% Kolliphor in 0.9% NaCl solution). For repetitive (4 days) HDAC inhibition during water maze training, Cl-994 was orally administered 1 h before the first training trial for four consecutive days. Habituation: Mice were trained to drink corn oil from plastic syringes one week (for 2 days) before the start of the experiment. A syringe (1 ml) filled with 120 µl corn oil (10% DMSO) was hung into the home cage, the entire solution was consumed voluntarily within 5 min. HDACi during WM training: A syringe filled with 120 µl of HDACi solution (0.8 mg Cl-994, 10% DMSO, 90% corn oil (Mazola)) or VEH solution (10% DMSO, 90% corn oil) was presented in front of the mouth until the entire solution was consumed.

### Behavioral experiments

For cFos re-expression experiments, TetTag mice from our colony were injected in two batches (batch 1: n = 2 mice for each condition; batch 2: n = 4 mice (HDACi), n = 3 mice (VEH)). Repetitive HDAC inhibition experiments were conducted in two batches of C57BL6/J mice (batch 1: n = 8 mice for each condition; batch 2: n = 4 mice (HDACi), n = 7 mice (VEH)). All experiments were recorded on digital video, Ethovision XT 17.5 was used for automated tracking. Animals were handled for 1 week before the start of the pre-training sessions to reduce stress during behavioral tasks. Pre-training: Mice were pre-trained for 1 day before their first exposure to the WM arena. Sessions (2 trials of max. 60 s) were done in a small rectangular water tank in the dark, in the same room where the WM task was performed. Water level was 1 cm above the escape platform (Ø 15 cm). Once the animals found the platform, a grid was presented until the animals climbed onto it and were returned to their home cages in the waiting area of the behavioral room. Training: The WM consisted of a circular tank (Ø 1.45 m) with visual asymmetrical landmarks, filled with water with (non-toxic) white paint. A platform (Ø 15 cm, submerged by 1 cm) was placed in the center of the target quadrant (first position, east; second position, west; third position, north) during training trails (max. swim time: 90 s). Before mice were trained to a new position, they were placed onto the new platform for 10 s to enhance reversal training. To test spatial reference memory, a probe trial (PT) without a platform was performed. PTs from day 1 until day 3 in the morning had a max. duration of 45 s (first 30 s are shown in the analysis, Fig. 7B,C; Fig. S7). After mice learned the second position, PT duration was increased to 60 s (Fig. 7E). After mice learned the third position, PT duration was further increased to 90 s (the first 60 s were analyzed, Fig. 7G). PTs on day 5 and day 11 had a duration of 120 s (the first 90 s were analyzed, Fig. 8). For the TTs, mice were lowered into the tank facing the wall in different, pseudo-randomized positions (avoiding the target quadrant). In PTs, mice were lowered in the center of the tank. An opaque cup attached to a pole was used to transfer the mice from their home cage to the drop position and a plastic grid attached to a pole was used to pick mice up. Mice were picked up 10 s after they found the platform and were returned to their home cage. Mice that did not find the platform during the TT were guided to it using the grid and were picked up after a 10 s on-platform waiting period. For both pre-training and WM, water temperature was 19 - 21°C and a heat lamp was placed over the waiting area to prevent hypothermia.

### Confocal imaging

To capture cFos and ΔFosB expression (Fig. 5) or cFos re-expression in the hippocampus during WM training (Fig. 6), immunostained slices from perfusion-fixed brains were imaged with a Zeiss LSM 900 (Plan-Apochromat, 20×/0.8). From each mouse, 4 - 6 slices (left and right DG, CA1) were imaged (stack of 5 image planes, 512 × 512 pixels, z-step: 3 µm) using the following filter sets: Alexa 405 (DAPI), Alexa 568 (mKate2, ΔFosB), Alexa 647 (cFos). The image acquisition settings were kept constant within an experimental data set. For display purposes, color channels were linearly adjusted in ImageJ.

### cFos re-expression analysis

Image stacks of the left and right hippocampus were analyzed with Imaris. The DAPI signal was used to create a volume for the upper and lower blade of the dentate gyrus. Based on reported cell densities (Jinno and Kosaka 2010), the volume was used to estimate the total number of granule cells. The cFos signal from both channels defined as cFos-tagged (568 nm) and 2^nd^ cFos ensemble (647 nm) were masked based on the DAPI volume. The cFos-tagged neurons were manually selected (8 µm spots) when all other channels were turned off. The 2^nd^ cFos ensemble was automatically detected using the quality filter function (see table 3). A Matlab based co-localization feature was used to identify overlapping spots from both populations (cFos-tagged, 2^nd^ cFos ensemble, closer than 5 µm). Based on the script the spots were assigned to cFos^+^ neurons, cFos-tagged neurons and double positive neurons. False positive spots were removed and non-detected spots were added. Ensemble size calculation and overlap analysis was done with Matlab. Lower cFos intensity spots (< 40) were removed and following parameters were calculated

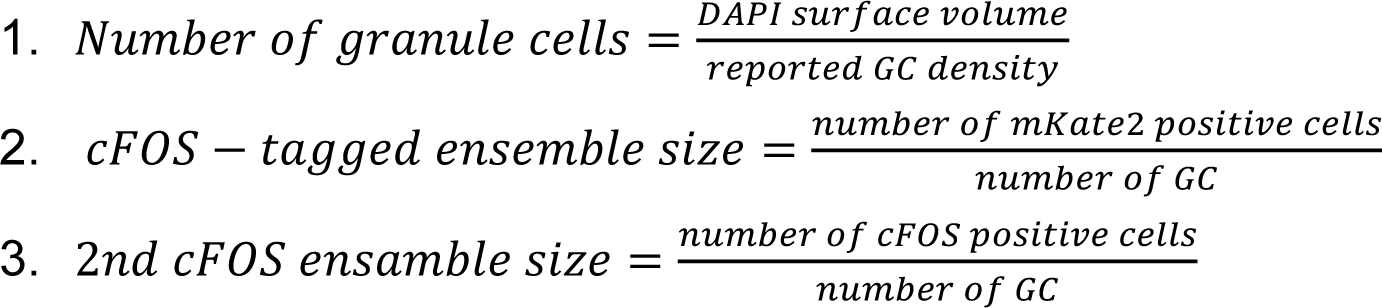

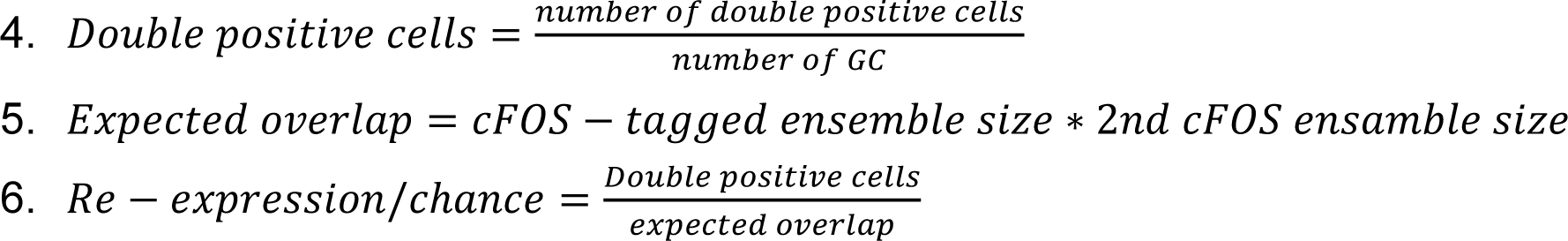

### Statistical analysis

Statistical tests were conducted in GraphPad Prism (version 9.01) and assumed an alpha level of 0.05. To analyze differences, we used one-way ANOVA, 2-way analysis of variance, and the Šídák method to correct for multiple comparisons. In some cases, we used two-sided t tests. A detailed description of experimental groups and statistical tests is provided as Supplementary Statistics. For all figures, *P < 0.05, **P < 0.01, ***P < 0.001, ****P < 0.0001.

## Acknowledgements

We thank Jan Schöder, Iris Ohmert and Michaela Schweizer for excellent technical assistance and Ingke Braren from the UKE vector facility for the production of rAAV. The ΔFosB plasmid was a gift from Eric Nestler (Addgene # 68545). The pAAV-TRE3G-BiPOLES-mKate2 construct (Addgene #192579) was a gift from J. Simon Wiegert.

## Author contributions

Conceptualization: T.G.O., F.M., C.E.G. and A.Fr. Experiments and data analysis: A.Fr. and P.J.L.M. Protein engineering: E.S. Fiber photometry: A.Fo. Writing— original draft: A.Fr. and T.G.O. Writing—review and editing: all authors.

## Data and materials availability

All data needed to evaluate the conclusions in the paper are present in the paper and/or the Supplementary Materials. Nuclear localized CaMPARI2 is available from the authors on request.

## References

1. Bading, Hilmar. 2013. “Nuclear Calcium Signalling in the Regulation of Brain Function.” Nature Reviews. Neuroscience 14 (9): 593–608.

2. Bakker, Arnold, C. Brock Kirwan, Michael Miller, and Craig E. L. Stark. 2008. “Pattern Separation in the Human Hippocampal CA3 and Dentate Gyrus.” Science (New York, N.Y.) 319 (5870): 1640–42.

3. Bengtson, C. Peter, H. Eckehard Freitag, Jan-Marek Weislogel, and Hilmar Bading. 2010. “Nuclear Calcium Sensors Reveal That Repetition of Trains of Synaptic Stimuli Boosts Nuclear Calcium Signaling in CA1 Pyramidal Neurons.” Biophysical Journal 99 (12): 4066–77.

4. Berron, D., H. Schutze, A. Maass, A. Cardenas-Blanco, H. J. Kuijf, D. Kumaran, and E. Duzel. 2016. “Strong Evidence for Pattern Separation in Human Dentate Gyrus.” The Journal of Neuroscience: The Official Journal of the Society for Neuroscience 36 (29): 7569–79.

5. Cates, Hannah M., Casey K. Lardner, Rosemary C. Bagot, Rachael L. Neve, and Eric J. Nestler. 2019. “Fosb Induction in Nucleus Accumbens by Cocaine Is Regulated by E2F3a.” ENeuro 6 (2): ENEURO.0325-18.2019.

6. Chatzi, Christina, Yingyu Zhang, Wiiliam D. Hendricks, Yang Chen, Eric Schnell, Richard H. Goodman, and Gary L. Westbrook. 2019. “Exercise-Induced Enhancement of Synaptic Function Triggered by the Inverse BAR Protein, Mtss1L.” ELife 8 (June). 10.7554/eLife.45920.

7. Chowdhury, Ananya, and Pico Caroni. 2018. “Time Units for Learning Involving Maintenance of System-Wide CFos Expression in Neuronal Assemblies.” Nature Communications 9 (1): 4122.

8. Coba, Marcelo P., Noboru H. Komiyama, Jess Nithianantharajah, Maksym V. Kopanitsa, Tim Indersmitten, Nathan G. Skene, Ellie J. Tuck, et al. 2012. “TNiK Is Required for Postsynaptic and Nuclear Signaling Pathways and Cognitive Function.” The Journal of Neuroscience: The Official Journal of the Society for Neuroscience 32 (40): 13987–99.

9. Creighton, Samantha D., Gilda Stefanelli, Anas Reda, and Iva B. Zovkic. 2020. “Epigenetic Mechanisms of Learning and Memory: Implications for Aging.” International Journal of Molecular Sciences 21 (18): 6918.

10. Eagle, Andrew L., Paula A. Gajewski, Miyoung Yang, Megan E. Kechner, Basma S. Al Masraf, Pamela J. Kennedy, Hongbing Wang, Michelle S. Mazei-Robison, and Alfred J. Robison. 2015. “Experience-Dependent Induction of Hippocampal ΔFosB Controls Learning.” The Journal of Neuroscience: The Official Journal of the Society for Neuroscience 35 (40): 13773– 83.

11. Eagle, Andrew L., Elizabeth S. Williams, Joseph A. Beatty, Charles L. Cox, and Alfred J. Robison. 2018. “ΔFosB Decreases Excitability of Dorsal Hippocampal CA1 Neurons.” ENeuro 5 (4): ENEURO.0104-18.2018.

12. Hainmueller, Thomas, and Marlene Bartos. 2018. “Parallel Emergence of Stable and Dynamic Memory Engrams in the Hippocampus.” Nature 558 (7709): 292–96.

13. Hevner, Robert F. 2016. “Evolution of the Mammalian Dentate Gyrus.” The Journal of Comparative Neurology 524 (3): 578–94.

14. Holbro, Niklaus, Asa Grunditz, and Thomas G. Oertner. 2009. “Differential Distribution of Endoplasmic Reticulum Controls Metabotropic Signaling and Plasticity at Hippocampal Synapses.” Proceedings of the National Academy of Sciences of the United States of America 106 (35): 15055–60.

15. Jaeger, Baptiste N., Sara B. Linker, Sarah L. Parylak, Jerika J. Barron, Iryna S. Gallina, Christian D. Saavedra, Conor Fitzpatrick, et al. 2018. “A Novel Environment-Evoked Transcriptional Signature Predicts Reactivity in Single Dentate Granule Neurons.” Nature Communications 9 (1): 3084.

16. Jinno, Shozo, and Toshio Kosaka. 2010. “Stereological Estimation of Numerical Densities of Glutamatergic Principal Neurons in the Mouse Hippocampus.” Hippocampus 20 (7): 829–40.

17. Johnston, Graham A. R. 2013. “Advantages of an Antagonist: Bicuculline and Other GABA Antagonists.” British Journal of Pharmacology 169 (2): 328–36.

18. Lamothe-Molina, Paul J., Andreas Franzelin, Lennart Beck, Dong Li, Lea Auksutat, Tim Fieblinger, Laura Laprell, et al. 2022. “ΔFosB Accumulation in Hippocampal Granule Cells Drives CFos Pattern Separation during Spatial Learning.” Nature Communications 13 (1): 6376.

19. Leutgeb, Jill K., Stefan Leutgeb, May-Britt Moser, and Edvard I. Moser. 2007. “Pattern Separation in the Dentate Gyrus and CA3 of the Hippocampus.” Science 315 (5814): 961–66.

20. Liu, Xu, Steve Ramirez, Petti T. Pang, Corey B. Puryear, Arvind Govindarajan, Karl Deisseroth, and Susumu Tonegawa. 2012. “Optogenetic Stimulation of a Hippocampal Engram Activates Fear Memory Recall.” Nature 484 (7394): 381–85.

21. Madar, Antoine D., Laura A. Ewell, and Mathew V. Jones. 2019. “Pattern Separation of Spiketrains in Hippocampal Neurons.” Scientific Reports 9 (1): 5282.

22. Maravall, M., Z. F. Mainen, B. L. Sabatini, and K. Svoboda. 2000. “Estimating Intracellular Calcium Concentrations and Buffering without Wavelength Ratioing.” Biophysical Journal 78 (5): 2655– 67.

23. Mellentin, C., H. Jahnsen, and W. C. Abraham. 2007. “Priming of Long-Term Potentiation Mediated by Ryanodine Receptor Activation in Rat Hippocampal Slices.” Neuropharmacology 52 (1): 118– 25.

24. Moeyaert, Benjamien, Graham Holt, Rajtarun Madangopal, Alberto Perez-Alvarez, Brenna C. Fearey, Nicholas F. Trojanowski, Julia Ledderose, et al. 2018. “Improved Methods for Marking Active Neuron Populations.” Nature Communications 9 (1): 4440.

25. Morellini, Fabio. 2013. “Spatial Memory Tasks in Rodents: What Do They Model?” Cell and Tissue Research 354 (1): 273–86.

26. Mozolewski, Pawel, Maciej Jeziorek, Christoph M. Schuster, Hilmar Bading, Bess Frost, and Radek Dobrowolski. 2021. “The Role of Nuclear Ca2+ in Maintaining Neuronal Homeostasis and Brain Health.” Journal of Cell Science 134 (8). 10.1242/jcs.254904.

27. Nestler, Eric J. 2014. “Epigenetic Mechanisms of Drug Addiction.” Neuropharmacology 76 (January): 259–68.

28. Neunuebel, Joshua P., and James J. Knierim. 2014. “CA3 Retrieves Coherent Representations from Degraded Input: Direct Evidence for CA3 Pattern Completion and Dentate Gyrus Pattern Separation.” Neuron 81 (2): 416–27.

29. Peixoto, Lucia, and Ted Abel. 2013. “The Role of Histone Acetylation in Memory Formation and Cognitive Impairments.” Neuropsychopharmacology: Official Publication of the American College of Neuropsychopharmacology 38 (1): 62–76.

30. Pettit, Noah L., Ee-Lynn Yap, Michael E. Greenberg, and Christopher D. Harvey. 2022. “Fos Ensembles Encode and Shape Stable Spatial Maps in the Hippocampus.” Nature 609 (7926): 327–34.

31. Preibisch, Stephan, Stephan Saalfeld, and Pavel Tomancak. 2009. “Globally Optimal Stitching of Tiled 3D Microscopic Image Acquisitions.” *Bioinformatics (Oxford*, England) 25 (11): 1463–65.

32. Ramamoorthi, Kartik, Robin Fropf, Gabriel M. Belfort, Helen L. Fitzmaurice, Ross M. McKinney, Rachael L. Neve, Tim Otto, and Yingxi Lin. 2011. “Npas4 Regulates a Transcriptional Program in CA3 Required for Contextual Memory Formation.” *Science (New York*, N.Y.) 334 (6063): 1669–75.

33. Ramirez-Mejia, Gerardo, Elvi Gil-Lievana, Oscar Urrego-Morales, Ernesto Soto-Reyes, and Federico Bermúdez-Rattoni. 2021. “Class I HDAC Inhibition Improves Object Recognition Memory Consolidation through BDNF/TrkB Pathway in a Time-Dependent Manner.” Neuropharmacology 187 (108493): 108493.

34. Reijmers, Leon G., Brian L. Perkins, Naoki Matsuo, and Mark Mayford. 2007. “Localization of a Stable Neural Correlate of Associative Memory.” *Science (New York*, N.Y.) 317 (5842): 1230–33.

35. Renthal, William, Tiffany L. Carle, Ian Maze, Herbert E. Covington 3rd, Hoang-Trang Truong, Imran Alibhai, Arvind Kumar, Rusty L. Montgomery, Eric N. Olson, and Eric J. Nestler. 2008. “Delta FosB Mediates Epigenetic Desensitization of the C-Fos Gene after Chronic Amphetamine Exposure.” The Journal of Neuroscience: The Official Journal of the Society for Neuroscience 28 (29): 7344–49.

36. Robison, Alfred J., and Eric J. Nestler. 2022. “ΔFOSB: A Potentially Druggable Master Orchestrator of Activity-Dependent Gene Expression.” ACS Chemical Neuroscience 13 (3): 296–307.

37. Ulery, Paula G., Gabby Rudenko, and Eric J. Nestler. 2006. “Regulation of DeltaFosB Stability by Phosphorylation.” The Journal of Neuroscience: The Official Journal of the Society for Neuroscience 26 (19): 5131–42.

38. Ulery-Reynolds, P. G., M. A. Castillo, V. Vialou, S. J. Russo, and E. J. Nestler. 2009. “Phosphorylation of DeltaFosB Mediates Its Stability in Vivo.” Neuroscience 158 (2): 369–72.

39. Vanhoutte, Peter, Jean-Vianney Barnier, Bernard Guibert, Christiane Pagès, Marie-Jo Besson, Robert A. Hipskind, and Jocelyne Caboche. 1999. “Glutamate Induces Phosphorylation of Elk-1 and CREB, along with c *-Fos* Activation, via an Extracellular Signal-Regulated Kinase-Dependent Pathway in Brain Slices.” Molecular and Cellular Biology 19 (1): 136–46.

40. Vecsey, Christopher G., Joshua D. Hawk, K. Matthew Lattal, Joel M. Stein, Sara A. Fabian, Michelle A. Attner, Sara M. Cabrera, et al. 2007. “Histone Deacetylase Inhibitors Enhance Memory and Synaptic Plasticity via CREB:CBP-Dependent Transcriptional Activation.” The Journal of Neuroscience: The Official Journal of the Society for Neuroscience 27 (23): 6128–40.

41. Villain, Hélène, Cédrick Florian, and Pascal Roullet. 2016. “HDAC Inhibition Promotes Both Initial Consolidation and Reconsolidation of Spatial Memory in Mice.” Scientific Reports 6 (1). 10.1038/srep27015.

42. Watrous, Andrew J., and Arne D. Ekstrom. 2014. “The Spectro-Contextual Encoding and Retrieval Theory of Episodic Memory.” Frontiers in Human Neuroscience 8 (February): 75.

43. Weng, Feng-Ju, Rodrigo I. Garcia, Stefano Lutzu, Karina Alviña, Yuxiang Zhang, Margaret Dushko, Taeyun Ku, et al. 2018. “Npas4 Is a Critical Regulator of Learning-Induced Plasticity at Mossy Fiber-CA3 Synapses during Contextual Memory Formation.” Neuron 97 (5): 1137–1152.e5.

44. Wiegert, J. Simon, Christine E. Gee, and Thomas G. Oertner. 2017. “Single-Cell Electroporation of Neurons.” Cold Spring Harbor Protocols 2017 (2): db.prot094904.

45. You, Jason C., Kavitha Muralidharan, Jin W. Park, Iraklis Petrof, Mark S. Pyfer, Brian F. Corbett, John J. LaFrancois, et al. 2017. “Epigenetic Suppression of Hippocampal Calbindin-D28k by ΔFosB Drives Seizure-Related Cognitive Deficits.” Nature Medicine 23 (11): 1377–83.

46. You, Jason C., Gabriel S. Stephens, Chia-Hsuan Fu, Xiaohong Zhang, Yin Liu, and Jeannie Chin. 2018. “Genome-Wide Profiling Reveals Functional Diversification of ΔFosB Gene Targets in the Hippocampus of an Alzheimer’s Disease Mouse Model.” PloS One 13 (2): e0192508.

