## Supplemental figures 1-8 for "Epigenetic repression of cFos supports sequential formation of distinct spatial memories"

### Supplementary Figures 1-8

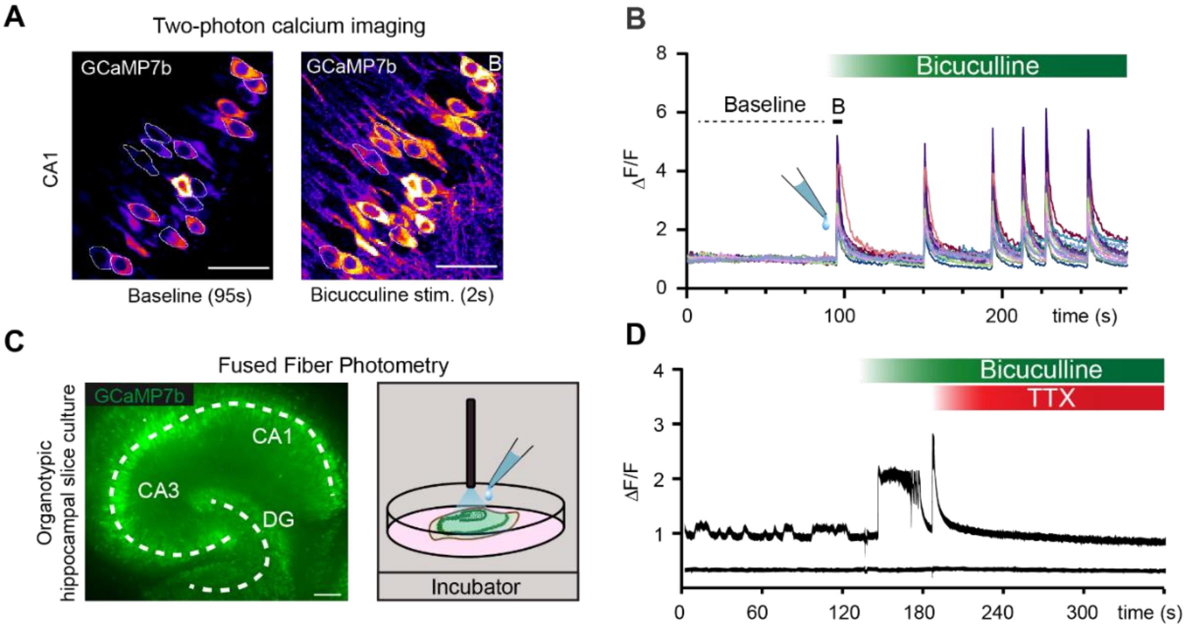


**Supplementary figure 1: Synchronized bursting after GABA$_A$ receptor block. A)** Organotypic hippocampal slice culture (rat) transduced with AAV9-Syn-jGCaMP7b (7x10$^{12}$ in buffer) (Dana et al. 2019). Left: GCaMP7b fluorescence at baseline, average of 190 images (256 x 256 pixels) acquired at 2 frames/s on a two-photon microscope at RT, peak-scaled to the brightest pixels. White lines around cell bodies show regions of interest (ROIs, n = 14). Right: Fluorescence after addition of bicuculline (20 µM), peak-scaled average of 4 images. Images were median-filtered (2-pixel radius), scale bars are 20 µm. **B)** GCaMP7b fluorescence intensity changes ($\Delta F/F_0$) relative to baseline fluorescence ($F_0$) in 14 neurons (ROIs shown in A) during bicuculline stimulation. Thick line 'B' indicates the first synchronous increase in jGCaMP7b intensity (right image in A). Six bursts of activity occurred within 200 s. **C)** Fused-fiber photometry (Formozov, Dieter, and Wiegert 2023) of jGCaMP7b fluorescence during bicuculline stimulation inside the incubator (37°C). Hippocampal slice cultures were virally transduced with jGCaMP7b and placed directly (1 mm) under the optical fiber inside the incubator. **D)** jGCaMP7b fluorescence intensity changes ($\Delta F/F_0$) during sequential application of bicuculline (20 µM, t = 140 s) and TTX (1 µM , t = 190 s) relative to baseline ($F_0$), 470 nm excitation. Note spontaneous activity at baseline, but not after TTX. The trace below the jGCaMP7b signal was recorded quasi-simultaneously with temporally interleaved 405 nm light pulses (isosbestic wavelength) to control for possible artifacts induced by application of bicuculline and TTX. Both functional and isosbestic signals were corrected by subtracting the autofluorescence of the optical fiber.

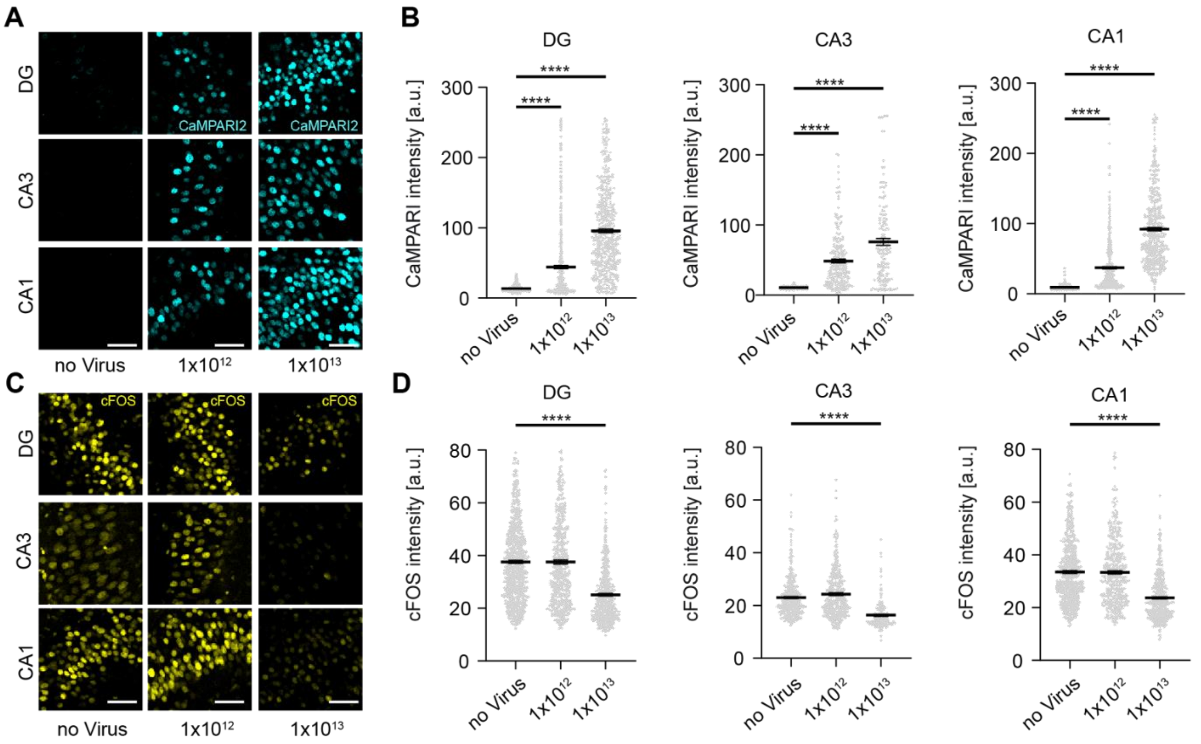


**Supplementary figure 2: Low H2B-CaMPARI2 level does not affect bicuculline-induced cFOS expression. A)** Confocal images of hippocampal subfields (DG, CA3, CA1) in slice cultures transfected with high ($10^{13}$ vg/ml) or low ($10^{12}$ vg/ml) titer of AAV.PhP.eb-syn-H2B-CaMPARI2 (cyan). Non-transfected cultures were used as controls. Scale bars = 20 µm. **B)** Single cell intensity analysis of H2B-CaMPARI2 expression (arbitrary units) in hippocampal subfields. For each condition, cells from 3 slice cultures were pooled. Gray dots show individual cells, black lines indicate mean ± SEM (DG, no virus, n = 585 cells; $10^{12}$, n = 428 cells, $10^{13}$, n = 427 cells; CA3, no virus, n = 704 cells; $10^{12}$, n = 510 cells, $10^{13}$, n = 468 cells; CA1, no virus, n = 370 cells; $10^{12}$, n = 233 cells, $10^{13}$, n = 158 cells). In all hippocampal subfields, expression level increased with virus titer (****p < 0.0001, $1x10^{13}$ vs no virus; ****p < 0.0001, $1x10^{12}$ vs no virus, 1-way-ANOVA, Dunett's multiple comparison). **C)** Bicuculline-induced cFOS expression (yellow), same regions as in panel A. Scale bars = 20 µm. **D)** Single cell intensity analysis of cFOS expression (same cells as in B). For all subfields, low virus titer ($1x10^{12}$) did not affect cFOS expression (DG, ns p = 0.99; CA3, ns p = 0.053; CA1, ns p = 0.97) whereas high virus titer ($1x10^{13}$) significantly reduced cFOS expression (DG, CA3, CA1, ****p < 0.0001, 1-way-ANOVA, Dunett's multiple comparison).

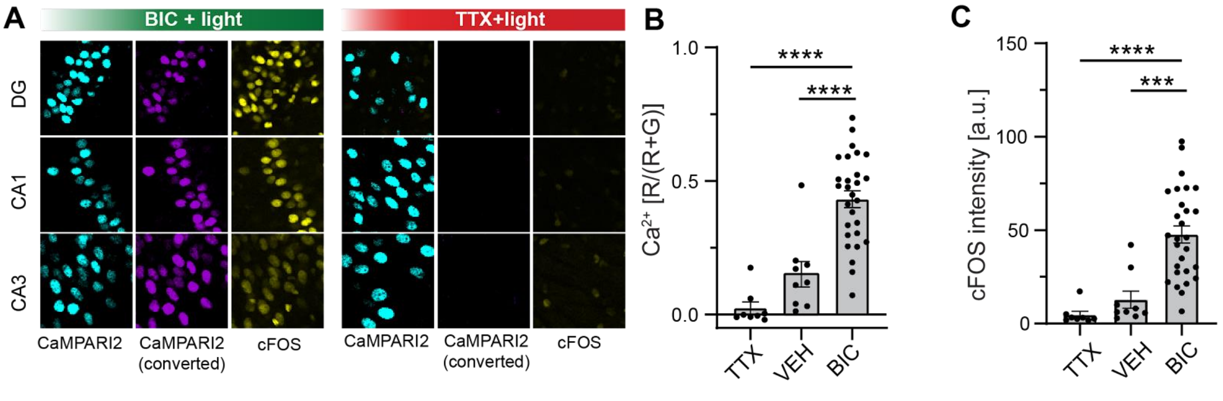


**Supplementary figure 3: No conversion of H2B-CaMPARI2 in the absence of activity. A)** Left: Confocal images of hippocampal subfields (DG, CA1, CA3) expressing H2B-CaMPARI2 after bicuculline stimulation (60 s, 20 µM, 100 µl in buffer) and continuous photoconversion with 395 nm light (green bar). Activity was terminated after 60 s by adding TTX (1 µM, 100 µl in buffer). Slice cultures were fixed 60 min later and stained for converted CaMPAIR2 (magenta) and cFOS (yellow). Green CaMPARI2 signal (cyan) was not immuno-enhanced. Right: Slice cultures treated with TTX (60 s, 1 µM, 100 µl) and illuminated for 60 s. **B)** Analysis of H2B-CaMPARI2 conversion (R/(R+G)) during 60 s of illumination after treatment with TTX, vehicle (VEH, 2.7% DMSO in buffer), or bicuculline. Based on the baseline fluorescence of CaMPARI2, 20 nuclei per subfield were selected for analysis (ROIs). Each black dot represents the mean of 20 ROIs, gray bars indicate mean of means ± SEM (TTX, n = 8; VEH, n = 9; BIC, n = 28). Photoconversion was significantly stronger after bicuculline stimulation compared to TTX (****$p < 0.0001$) or vehicle-treated cultures (****$p < 0.0001$). Th difference between VEH and TTX conditions was not significant (ns, $p = 0.20$, 1-way-ANOVA, Dunett's multiple comparison). **C)** cFOS intensity analysis, same ROIs as in B). cFOS levels in bicuculline-stimulated slice cultures were significantly higher compared to TTX (****$p < 0.0001$) or vehicle-treated cultures (***$p = 0.0002$). The difference between VEH and TTX conditions was not significant ($p = 0.69$, 1-way-ANOVA, Dunett's multiple comparison). Note: BIC data are identical to the one-time stimulated cultures presented in Fig. 4.

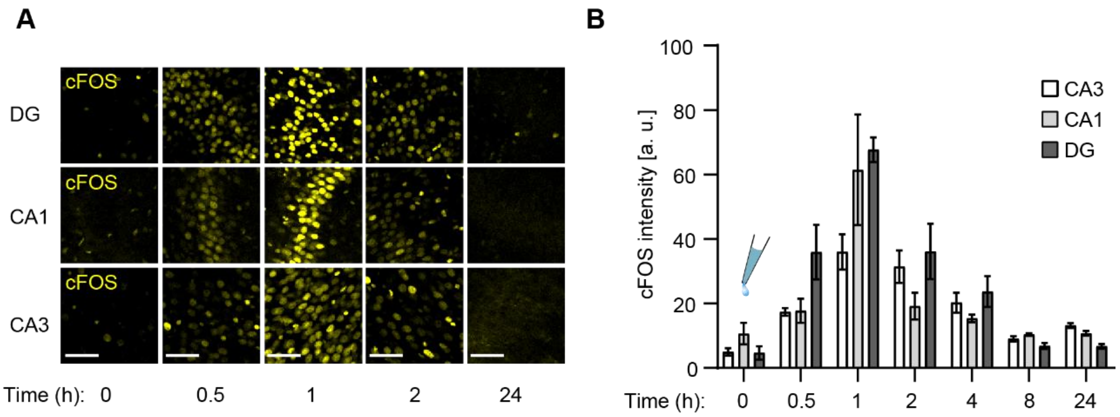


**Supplementary figure 4: Temporal profile of cFOS expression after bicuculline stimulation at t = 0.**
**A)** Confocal images of hippocampal subfields (DG, CA1, CA3) stimulated with bicuculline (100 µl, 20 µM in buffer). Stimulation was terminated after 60 s by adding TTX (tetrodotoxin, 300 µl, 1 µM in buffer). Slice cultures were fixed after t = 0, 0.5, 1, 2 or 24 hours and stained for cFOS (yellow) and DAPI (not displayed). Scale bar = 50 µm. **B)** cFOS intensity analysis (arbitrary units). Bars (white = CA3, gray = CA1, dark gray = DG) represent mean ± SEM from n = 3 slice cultures per time point. From each hippocampal subfield, 20 ROIs were randomly selected based on the DAPI signal. During ROI selection, the cFOS channel was turned off. In all subfields, cFOS intensity peaked 1 h after bicuculline stimulation.

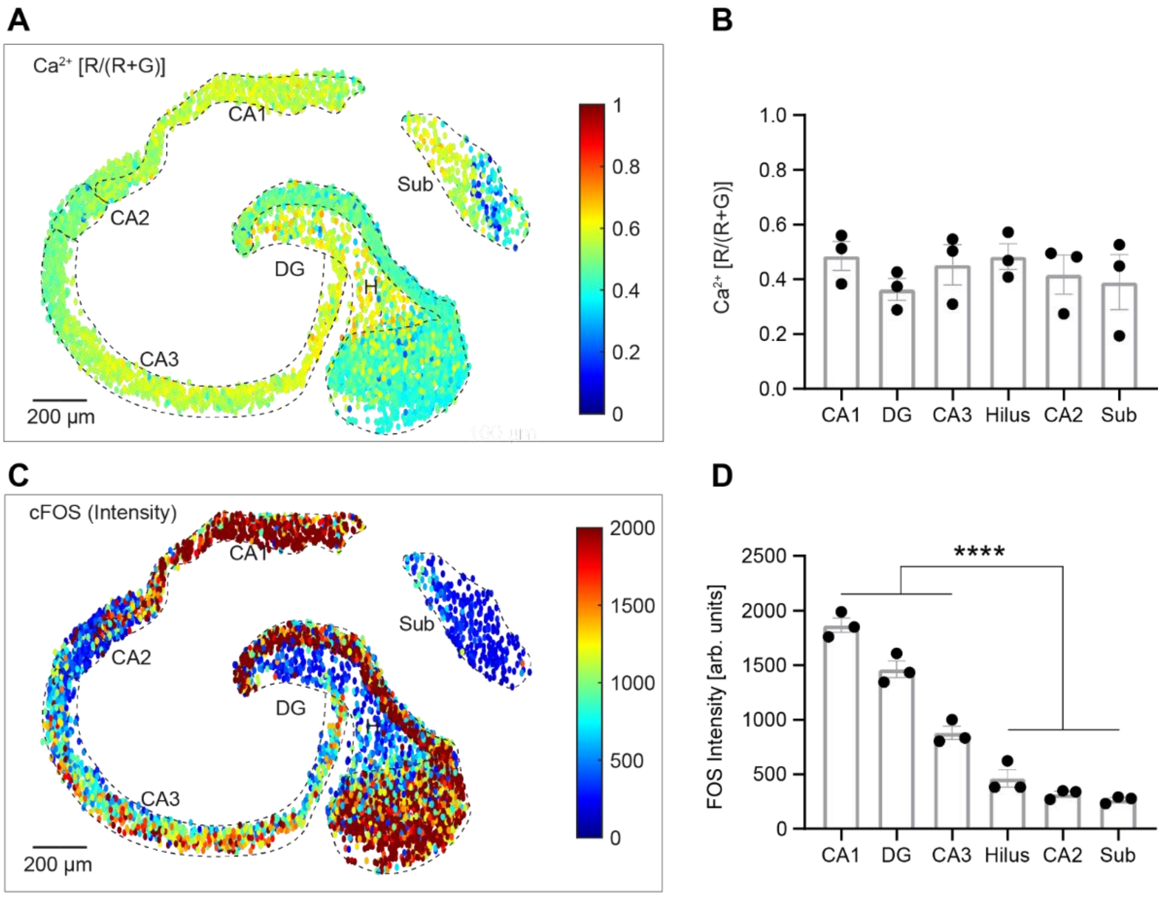


**Supplementary figure 5: Spatial analysis of $Ca^{2+}$ and cFOS expression after bicuculline stimulation.**
**A)** H2B-CaMPARI2-positive nuclei were automatically detected using Imaris (Spot size: 8 µm). For each spot, the conversion ratio (R/(R+G)), corresponding to nuclear $Ca^{2+}$ levels, was calculated in Matlab. Spots were color-coded according to the normalized conversion ratio (ranging from 0 to 1) and plotted at their spatial coordinates (CA1, n = 606 nuclei; CA2, n = 247 nuclei; CA3, n = 830 nuclei; DG, n = 2245 nuclei; hilus (H), n = 294 nuclei; subiculum (Sub), n = 343 nuclei). Scale bar = 200 µm. **B)** Analysis of mean $Ca^{2+}$ level ± SEM per region (n = 3 slice cultures). Neurons of all regions showed similar $Ca^{2+}$ levels. **C)** Same analysis procedure like in panel A, but spots are color-coded based on their cFOS intensity (0-2000, arbitrary units). Scale bar = 200 µm. **D)** Analysis of mean cFOS level ± SEM per region (n = 3 slice cultures). Principal neurons in CA1, CA3 and DG show significantly higher cFOS expression compared to neurons in CA2, the hilus and the subiculum (****$p < 0.0001$; t-test).

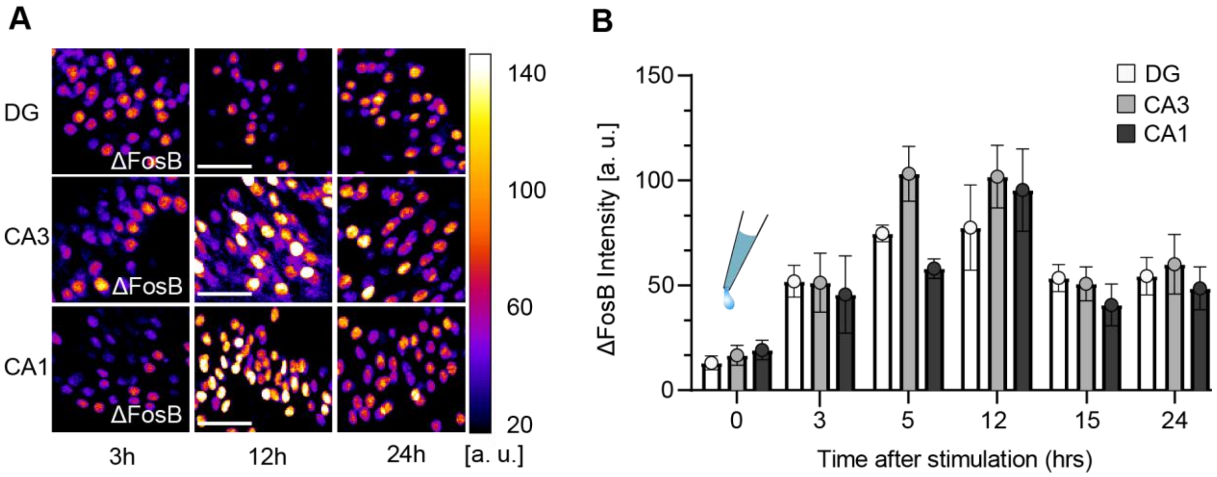


**Supplementary figure 6: Temporal profile of ΔFosB expression after bicuculline stimulation at t = 0.**
**A)** Confocal images of hippocampal subfields (DG, CA1, CA3) stimulated with bicuculline (100 µl, 20 µM in buffer). Stimulation was terminated after 60 s by adding TTX (300 µl, 1 µM in buffer). Slice cultures were fixed after t = 0, 3, 5, 12, 15 and 24 hours and stained for ΔFosB (fire color scale, arbitrary units) and DAPI (not displayed). Scale bar = 50 µm. B) ΔFosB intensity analysis. Bars (white = DG, gray = CA3, dark gray = CA1) represent mean ± SEM from 3 - 8 slice cultures per time point. ΔFosB intensity peaked 5 -12 hours after bicuculline stimulation and was still elevated after 24 h.

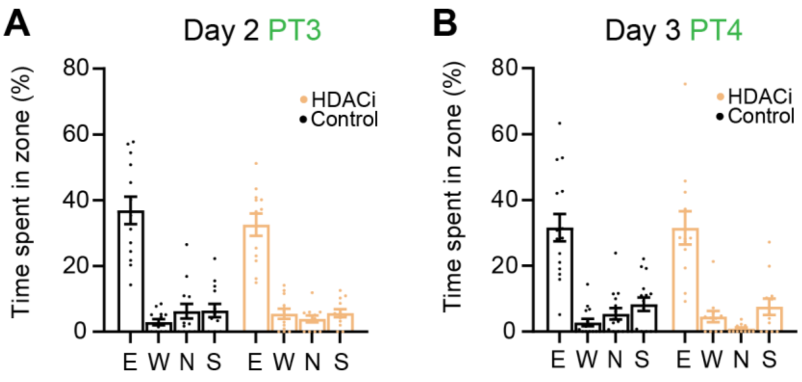


**Supplementary figure 7: Spatial memory recall on day 2 and day 3. A)** Third probe trail for platform position east (E, first trained position) on day 2. Each dot represents one animal, bars show mean ± SEM. Both groups (VEH, black, n = 14; HDACi, ocher, n = 12) show the same preference for the trained target zone (E) during the first 30 s of a 45-second probe trail (E, east, p = 0.2; W, west, p = 0.91; N, north, p = 0.97; S, south, p = 0.99). **B)** Fourth probe trail for platform position east (E, first position) on day 3. Each dot represents one animal, bars show mean ± SEM. Both groups (VEH, black, n = 15; HDACi, ocher, n = 12) show the same preference for the trained target zone (E) during the first 30 s of a 45-second probe trail (E, p > 0.99; W, p = 0.98; N, p = 0.68; S, p = 0.99, 2-way ANOVA, Šídák's multiple comparisons).

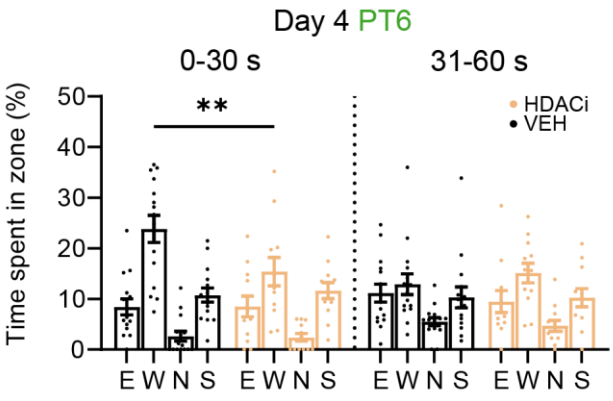


**Supplementary figure 8: Spatial memory recall on day 4.** Second probe trail (60 s) in the morning of day 4, last trained platform position west (W). Each dot represents one animal, bars show mean ± SEM. Behavior during the first and second half of the probe trial was analyzed separately (dotted line). During the first 30 s, vehicle-treated mice (n = 15) spent significantly more time around the last trained platform position (W) compared the HDACi group (n = 12) (E, east, p > 0.99; W, west, ** p = 0.0073; N, north, p > 0.99; S, south, p = 0.99). No difference in position preference was observed during the last 30 s between both conditions (E, p = 0.94; W, p = 0.86; N, p = 0.99; S, p > 0.99; 2-way ANOVA, Šídák's multiple comparison).

#### Supplementary References

Dana, Hod, Yi Sun, Boaz Mohar, Brad K. Hulse, Aaron M. Kerlin, Jeremy P. Hasseman, Getahun Tsegaye, et al. 2019. “High-Performance Calcium Sensors for Imaging Activity in Neuronal Populations and Microcompartments.” *Nature Methods* 16 (7): 649–57.

Formozov, Andrey, Alexander Dieter, and J. Simon Wiegert. 2023. “A Flexible and Versatile System for Multi-Color Fiber Photometry and Optogenetic Manipulation.” *Cell Reports Methods* 3 (3): 100418.
